# Cooperativity between Ras pathway mutations in colonic tumorigenesis

**DOI:** 10.1101/2025.09.05.673994

**Authors:** Christian W. Johnson, Anna Idelevich, Bing Shui, Alison Tisdale, Eve O’Donoghue, Josh Cook, Shannon Hull, Moon-Hee Yang, Shikha Sheth, Elizabeth M. Terrell, Stefania Bovo Minto, Deborah K. Morrison, Scott Kopetz, Rona Yaeger, Kevin M. Haigis

## Abstract

The Kirsten rat sarcoma (*KRAS*) gene is the most frequently mutated oncogene in colorectal cancer (CRC). We previously characterized two activating alleles of *KRAS*, A59T and A59E, that show impaired BRAF dimerization. These alleles of *KRAS* are enriched in CRC tumors with genetic alterations in genes that regulate MAPK signaling (e.g. EGFR, NF1). Using a new conditional mouse model of K-Ras (LSL-K-Ras A59E) we show that, despite its inability to universally activate RAF kinases, K-Ras A59E disturbs colon epithelium and decreases mouse survival in a tumor model. By combining LSL-K-Ras A59E mice with a conditional knockout of Nf1, we demonstrate cooperation between these alleles at multiple levels. Consistent with cooperation and clinical observations, we show that K-Ras A59E neither promotes EGF independence of organoid growth, nor confers intrinsic resistance to EGFR inhibition. Thus, our data provide a deeper understanding of the role of RAF isoforms in mutant KRAS driven CRC and argue for the use of anti-EGFR therapies against some Ala59 mutant alleles of KRAS.

**Statement of significance:** Our studies on oncogenic K-RAS mutants demonstrate that the biochemical properties of a mutant oncoprotein are reflected in the somatic genetics of cancer, in particular in the cooperating mutations that occur. These mutant-specific genetic interactions influence therapeutic responses. As such, *KRAS* mutation may not be a univariate predictor of response to EGFR inhibition.

## Introduction

Colorectal cancer (CRC) is a leading cause of cancer mortality in the US (1). The epidermal growth factor receptor (EGFR) inhibitors panitumumab and cetuximab (collectively EGFRi) can extend the survival of patients with metastatic CRC (mCRC) (1). This survival benefit is dependent on the tumor genotype and approximately 40% of mCRC patients are not treated with EGFRi due to the presence of *KRAS* mutations, which confer resistance because of presumed EGFR-independent activation of the MAPK signaling pathway (1). Nevertheless, mutant *KRAS* alleles are heterogenous, especially in CRC, and intrinsic resistance to EGFRi may not be a universal feature of oncogenic K-RAS (2). For example, mCRCs expressing the rare mutant K-RAS^A59T^ exhibit varied clinical responses to EGFRi, some of which are remarkable for cancers with a *KRAS* mutation (3,4). The mechanistic basis for varied EGFRi responses in KRAS-mutant CRC remains unclear.

Activation of the MAPK signaling pathway is the most important oncogenic function of K-RAS (5). Under normal physiological conditions, K-RAS is maintained in an inactive, GDP bound state by GTPase activating proteins (GAPs), such as NF1, which bind the active site of K-RAS and catalyze GTP hydrolysis. Upon upstream activation of a receptor tyrosine kinase (RTK), such as EGFR, by growth factors, K-RAS is recruited to the cell membrane and activated by guanine nucleotide exchange factors (GEFs). GEFs catalyze the exchange of GDP for GTP in the active site of K-RAS, which in turn allows it to recruit cytosolic RAF kinases to the membrane (6). Once at the membrane, monomeric RAF bound to K-RAS undergoes a conformational restructuring, re-organization of phosphorylation, and dimerization with other RAF proteins (6). Once dimerized, RAF proteins are fully activated and phosphorylate the dual-specificity kinases MEK1 and MEK2. MEK1/2 phosphorylate ERK1 and ERK2, which in turn translocate to the nucleus to promote changes in gene expression. Mutant forms of K-RAS favor the activated GTP-bound state (7) and, as a result, promote hyperactivation of the MAPK signaling pathway through stabilization of RAF heterodimers (8,9).

K-RAS mutations can be classed by their mechanism of activation and location in the three-dimensional structure of the protein (7). Class 1 mutations occur at Gly12 mutations and function by inhibiting GAP-mediated, and intrinsic GTP hydrolysis (10). Since class 1 mutations are the most frequent (2/3 of all *KRAS* mutations in CRC), insensitivity to GAP is considered a main determinant of K-RAS oncogenicity. GAPs have a more complex relationship with class 2 K-RAS mutants, such as those occurring at Ala146, Lys117, and Gly13. Class 2 mutations show variable sensitivity towards GAP proteins. For instance, G13D retains sensitivity to NF1, but not RASA1 (11), while A146T retains sensitivity to RASA1, and likely NF1, due to the location of this mutation within the active site of K-RAS (12). Class 2 mutations promote functional activation of K-RAS by negatively interacting with bound guanine nucleotides to promote an increase in nucleotide exchange. Since GTP is in excess in the cell, class 2 mutations promote a net increase in GTP-bound K-RAS. Finally, class 3 mutations, such as those at Gln61 and Ala59, also rely on enhanced nucleotide exchange, but with a more impaired capacity to promote GTP hydrolysis. Class 3 mutants also appear to influence the ability of K-RAS to interact with different effectors, including RAF kinases (7).

We recently described the mechanisms of K-RAS activation by A59T, its autophosphorylation at Thr59, and its phosphomimetic A59E (13). Unlike class 1 and 2 mutants of K-RAS, A59T and A59E mutations stabilize the active GTP-bound state of K-RAS by simultaneously inhibiting GTP hydrolysis and promoting nucleotide exchange (13–15). At the atomic level, this is due to A59T and A59E destabilizing the active site of K-RAS (13). Paradoxically, A59T and A59E are weak oncogenic alleles. This results from their reduced affinity for the Ras Binding Domain (RBD) of A-RAF, B-RAF, and C-RAF, resulting in an attenuated ability to recruit B-RAF to the cell membrane and promote B-RAF/C-RAF heterodimerization (13). Furthermore, biochemical studies have demonstrated that the A59E mutation, while of allowing K-RAS to associate with B-RAF or C-RAF, prevents activation of their kinase activities (16,17). Thus, A59T and A59E are partial gain-of-function mutants of K-RAS, capable of activating only a subset of signaling functions normally associated with oncogenic K-RAS. Here, we sought to understand how K-RAS mutants with attenuated oncogenicity actually contribute to CRC.

## Results

### A59E is a weak oncogenic allele of KRAS

Residue 59 mutants of K-RAS and H-RAS have been tested previously in cellular assays of oncogenic transformation (13,18). Because classical transformation assays typically take advantage of hyper-sensitivity of certain cell types (*e.g.* NIH3t3) to RAS activation, however, these assays do not effectively model how somatic mutations in endogenous K-RAS contribute to tumorigenesis in the gastrointestinal tract (12). We used CRISPR to genetically engineer a conditional allele of *Kras* under the control of a lox-STOP-lox cassette (LSL-K-Ras) in mouse embryonic stem cells to generate a Cre-dependent allele of K-Ras^A59E^ (i.e. *Kras*^LSL-A59E^) (**Supplementary Fig. 1A-C**). To study this allele in the mouse colon, we crossed *Kras*^LSL-A59E^ to *Fabpl1-Cre* mice, which express Cre in the epithelium of the distal small intestine (SI), cecum, and colon (19). Using these animals, and comparing them to mice carrying *Kras*^LSL-G12D^, we found that *Fabpl1-Cre*; *Kras*^LSL-A59E^ colons express normal levels of total K-Ras, but relatively low levels of K-Ras that can bind to the C-Raf RBD (**Fig. 1A**).

**Figure 1.**
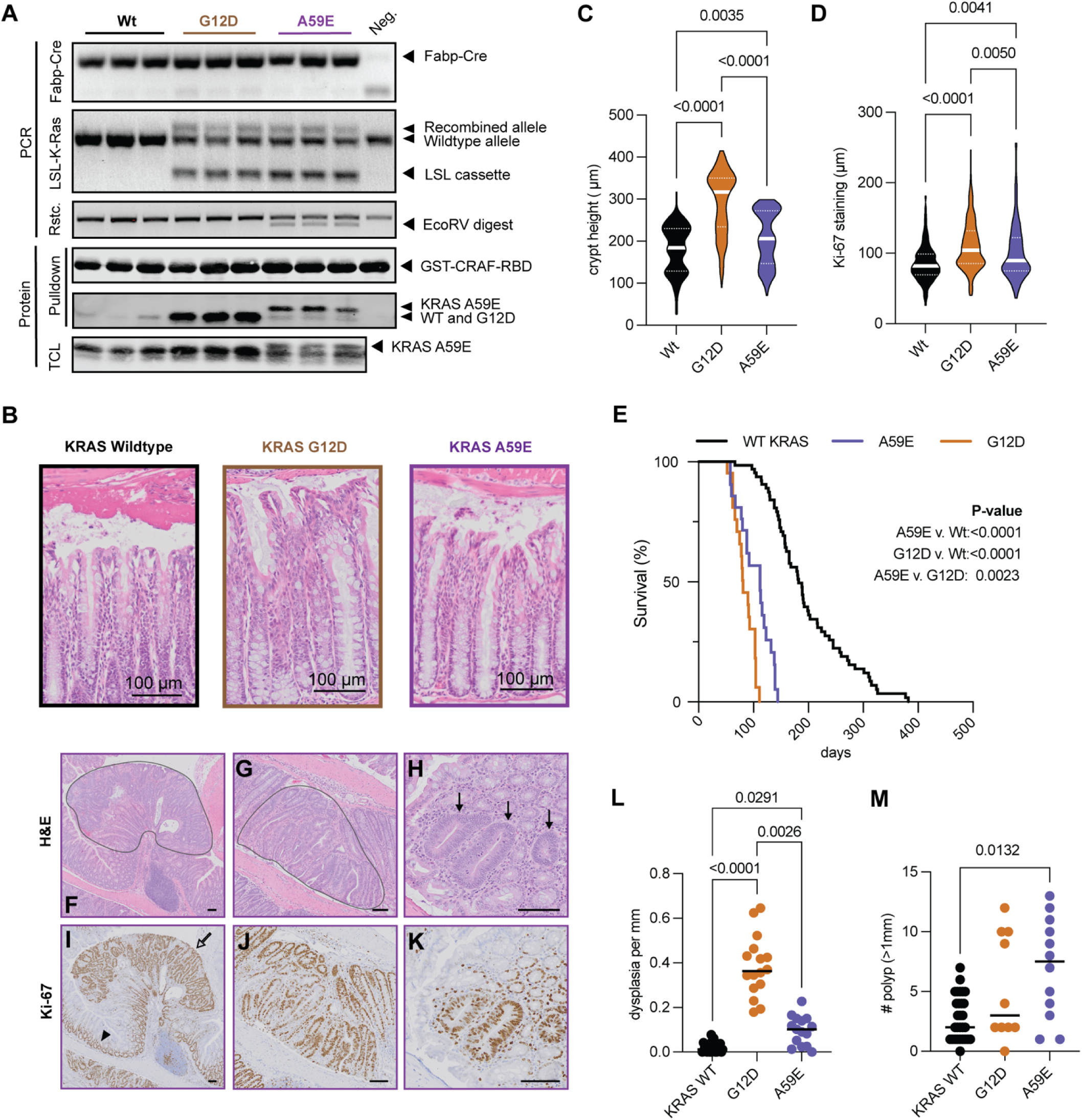
KRAS A59E is a weak oncogenic allele of the GI tract. (A) Validation of LSL-K-Ras A59E mouse. (B) H&E stain of mouse colon epithelium. (C) Quantification of crypt lengths. (D) Quantification of crypt proliferative zone by Ki-67 staining. (E) Survival curve of Fabp-cre(+), Apc(+/2lox14) mice either wildtype for KRAS (N=60), or harboring LSL-K-Ras G12D (N=19) or LSL-K-Ras A59E (N=20). P-values are the product of Log-Rank Mantel Cox tests for survival. (F-H) Representative H&E stains of different forms of dysplasia found in mice of genotype Fabp-Cre(+), Apc(+/2lox14), K-Ras(+/LSL-A59E). (F) Pedunculated polyp. (G,H) Sessile polyps. Polyps are outlined in grey (F,G) or by black arrows (H). Scale bars are 100µM. (I-K) Ki-67 stains of representative images from (F-H). Normal proliferative zone is indicated by the arrowhead and the polyp is indicated by the hollow arrow (I). Scale bars are 100µM. (L) Quantitation of dysplasia shown in (F-K). (M) Polyps greater than 1mm seen in different tumor mice. Statistical analysis of panels (L-M) was performed using Kruskal-Wallis tests with significance cutoff at <0.05. In all panels, * P < 0.05, ** P < 0.01, *** P < 0.001

When expressed in the colonic epithelium, mutant K-Ras does not promote neoplasia, but rather promotes crypt hyperplasia that is dependent on canonical MAPK pathway activation (12,20,21). In colons expressing K-Ras^A59E^ mutant mice, we observed a hyperplasia phenotype that was severely attenuated relative to that in colons expressing K-Ras^G12D^ (**Fig. 1B-D and Supplementary Fig. S1D,E**, suggesting that K-Ras^A59E^ can promote MAPK activation, but at a much lower level that K-Ras^G12D^. Consistent with this observation, we noted that K-Ras^A59E^ failed to induce colon lengthening as does K-Ras^G12D^ (**Supplementary Fig. S1F**).

In human and mouse colonic tumorigenesis, loss of the Adenomatous Polyposis Coli (APC) tumor suppressor gene acts as an initiating event and is found in >80% of CRC (22). Therefore, we next tested whether K-Ras^A59E^ can cooperate with APC loss, as we have previously demonstrated for K-Ras^G12D^ and K-Ras^A146T^ (12,20). In the context of the tumor model, where all mice carry a single *Apc*^2lox14^ floxed allele, K-Ras^A59E^ significantly reduced survival, but to a lesser extent than mice expressing K-Ras^G12D^ (**Fig. 1E**). At the morphological level, K-Ras^G12D^ cooperates with APC loss to promote dysplastic foci ranging from dysplastic aberrant crypt foci to sessile and pedunculated adenocarcinoma of the colon (12,20). Colons expressing K-Ras^A59E^ exhibited a similar range of colonic dysplasia compared to those expressing K-Ras^G12D^, including crypt disorganization, epithelial thickening, increase in the nuclear to cytoplasmic ratio (**Fig. 1F-H**), extension of proliferation past the normal proliferative zone (**Fig. 1I,J**), and nuclear stratification (**Fig. 1K**). Not surprisingly, K-Ras^G12D^ was better at promoting tumor formation when compared to K-Ras^A59E^ (**Fig. 1L**), however dysplastic lesions were significantly greater in size when expressing K-Ras^A59E^ (**Fig. 1M**). Thus, consistent with our observations in Apc wild-type colon, our data from colonic tumors indicate K-Ras^A59E^ and K-Ras^G12D^ induce quantitatively different cellular phenotypes.

### Ala59 mutants maintain MAPK signaling via A-RAF/C-RAF dimerization

We and others have shown that Ala59 mutations activate K-RAS while restricting B-RAF activation (13,16,17). Nevertheless, Ala59 mutations are still capable of promoting upregulation of MEK and ERK phosphorylation in mouse embryonic fibroblasts (13). Based on analysis of mice expressing K-Ras^A59E^ in the colonic epithelium, we reasoned that the intermediate hyperplasia phenotype induced by K-Ras^A59E^ was likely due to deficient levels of MAPK pathway activation. As expected, Mek and Erk phosphorylation in the colonic epithelia expressing K-Ras^A59E^ was intermediate in comparison to colons expressing WT or K-Ras^G12D^ (**Fig. 2A**), albeit not significantly different from either at a statistical level. Our prior signaling studies of K-Ras^A59E^ in mouse fibroblasts demonstrated that this mutant was deficient in promoting AKT phosphorylation (13). Indeed, when we examined AKT phosphorylation at Thr308 – phosphorylated by PDK1 – and Ser473 – phosphorylated by mTORC2 – in hyperplastic epithelium, we found no statistical difference between colon epithelia of the different genotypes (**Supplementary Fig. S1G**).

**Figure 2.**
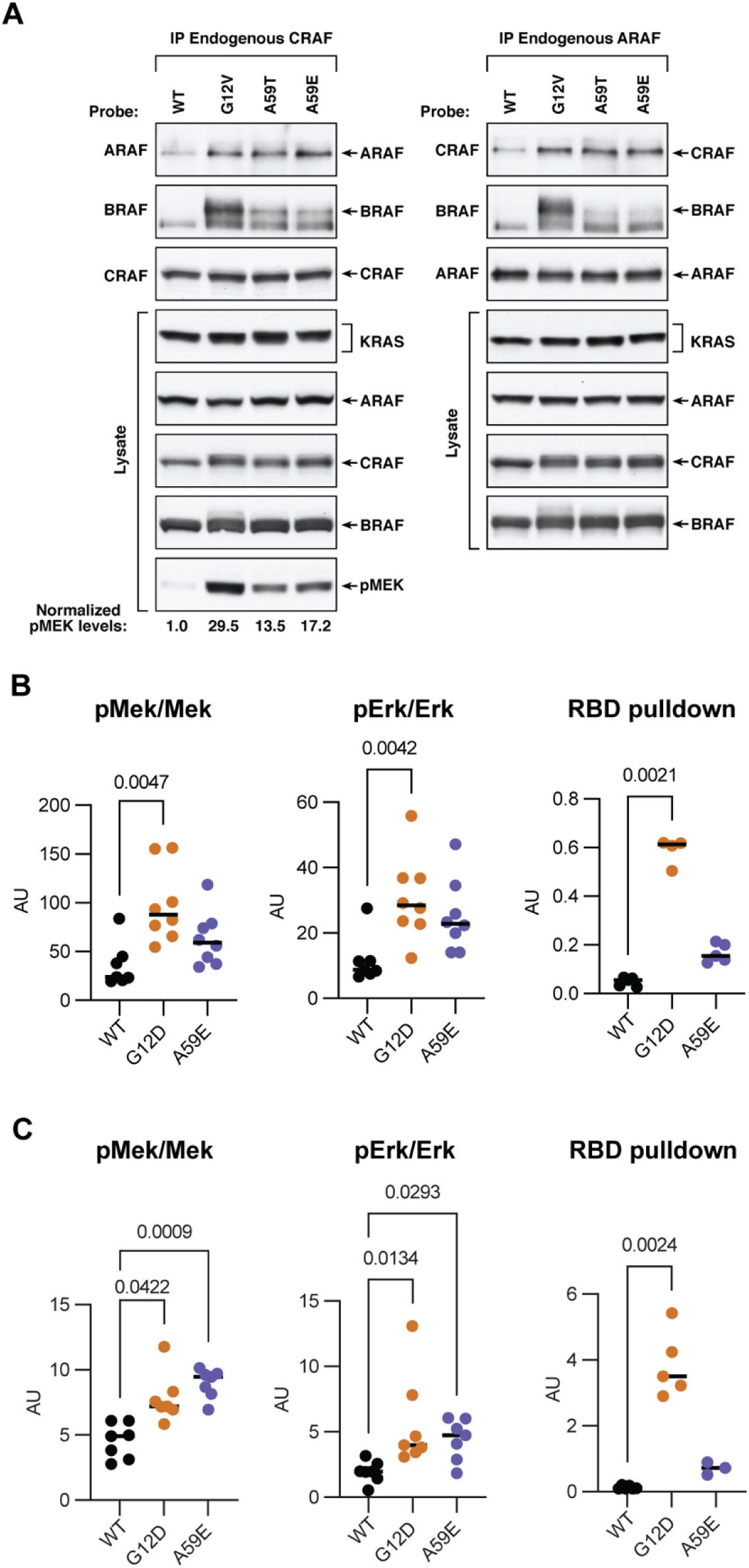
RAS pathway signaling in K-Ras A59E tissues and cells. (A) Phosphorylation of Mek (left) and Erk (middle) in colonic epithelium and tumors. RAF-RBD pulldown of Ras is shown on the right. (B) Co-immunoprecipitation of ARAF (left) of CRAF (right) for different RAF isoforms in HeLa cells expressing different KRAS mutants.

We then re-examined the ability of Ala59 mutants to promote RAF heterodimerization in human epithelial cells after ectopic expression of mutant K-Ras and discovered that A-RAF and C-RAF dimerization was unaffected by these mutations, while B-RAF/C-RAF and B-RAF/C-RAF dimerization was considerably attenuated in cells expressing K-Ras^A159T/E^ relative to K-Ras^G12V^ (**Fig. 2B**). Moreover, MEK phosphorylation in these cells was quantitatively lower for K-Ras^A59T/E^ compared with K-Ras^G12V^ (**Fig. 2B**). These data indicate that residue 59 K-Ras mutants exhibit both qualitative and quantitative differences in their ability to activate MAPK pathway signaling as compared to Gly12 mutants of K-Ras.

Finally, we examined colonic polyps for levels of activated Ras and for Mek and Erk phosphorylation. The relative activation states of K-Ras^G12D^ and K-Ras^A59E^ were the same between colon epithelia and pedunculated tumors (**Fig. 2C**). In contrast, K-Ras A59E induced levels of Mek and Erk phosphorylation comparable to K-Ras G12D (**Fig. 2C**). While it is unclear how Apc inactivation potentiates oncogenic K-Ras induced Mek/Erk phosphorylation, our data indicate that levels of Mek/Erk phosphorylation do not correlate with tumors phenotypes, such as overall survival and tumor number (**Fig. 1E-M**).

### Co-mutation patterns in CRC expressing Ala59 mutations of KRAS

K-Ras^A59T/E^ are examples of the complexity of the GTPase cycle and its relationship to oncogenic activity, as the mutations promote the GTP state by reducing GAP-induced hydrolysis, but also restrict oncogenic function by limiting interaction and activation with effectors, like B-RAF (13,16,17). To extend beyond our initial biochemical studies, we examined how the qualitative and quantitative differences in the activation of the MAPK signaling pathway by K-Ras^A59T/E^ could potentially influence mutational patterns in primary human cancers. We previously examined co-mutations of common and rare *KRAS* alleles in cancer and found that the cooperating mutations that occur in *KRAS* mutant cancer are allele- and tissue-specific (23,24). Using this original dataset and honing in on RTK/RAS pathway, as defined by The Cancer Genome Atlas (TCGA) (22), we characterized the frequency of mutations in pathway genes as a function of which K-RAS mutant is expressed in a given cancer. Colorectal cancers expressing K-RAS^A59T^ or K-RAS^A59E^ demonstrated a strong propensity for additional mutations in genes of the RTK/RAS pathway, more so than any other K-RAS mutant (**Fig. 3A**). As the dataset from Cook et al. was small (N=1536) with relatively few KRAS^A59T/E^ alleles (N=<10) (23), we next performed a meta-analysis of human tumors by combining published exome and whole genome sequencing data from TCGA and AACR GENIE, and unpublished data from Memorial Sloan Kettering and MD Anderson Cancer Center (**Supplementary Table S1**). The combined data confirmed that Ala59 mutations are enriched in cancers of gastrointestinal (GI) and hematological origin, and that A59T is the most common missense mutation overall (**Supplementary Fig. S2A-C**). However, while A59T and A59E are enriched in cancers of the gastrointestinal tract, we noted that A59G mutations predominate in hematological cancers (**Supplementary Fig. S2C**). Consistent with our initial analysis (**Fig. 3A**), GI tumors expressing K-RAS A59T or A59E were enriched for additional mutations in the RTK/RAS pathway, including genetic lesions in common oncogenes and tumor suppressors that are normally not found with mutant KRAS (**Fig. 3B**). Of note, cancers expressing K-RAS^A59G^ did not exhibit an enrichment in RTK/RAS co-mutations. This observation is consistent with the hypothesis that cancer genotype is related to the biochemical properties of activated oncogenes, as K-Ras^A59G^ is biochemically distinct from K-Ras^A59T/E^ (13,25).

**Figure 3.**
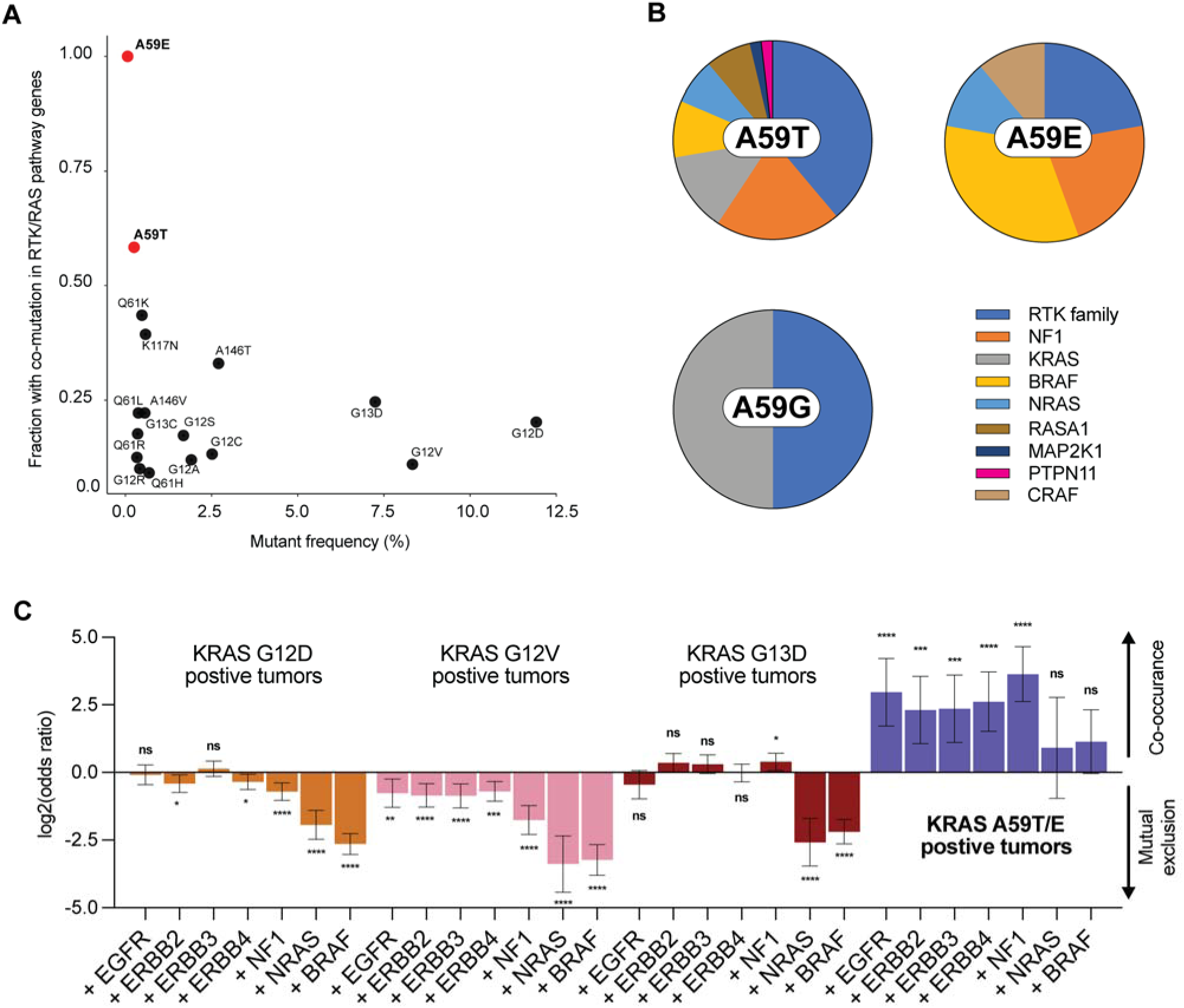
Ala59 mutants of KRAS favor additional activation of the RTK/RAS pathway in cancer. (A) Frequency of MAPK co-mutation in KRAS mutants using meta-dataset from Cook et al. 2020. (B) Frequency of driver RTK/RAS co-mutations in GI tumors harboring KRAS A59T (N=79), A59E (N=16), and A59G (N=4) alleles. See Supplementary Table 1. (C) Odds Ratios (y-axis) comparing likelihood of the indicated co-mutation (x-axis) in colorectal tumors positive for KRAS G12D, G12V, G13D, or KRAS A59T and A59E mutations. Statistics indicate two-sided Chi-square tests with Yates correction (* P-value <0.05, ** P-value <0.005, *** P-value <0.0005, **** P-value <0.0001). Error bars represent 95% confidence intervals calculated using the Gart adjusted logit interval. In bold are RTK family members.

We next calculated odds ratios (*i.e.* favorability of co-occurrence) for co-mutations in NRAS, NF1, BRAF, and RTK family genes with A59T or A59E KRAS mutants using the AACR GENIE dataset (18,334 samples) and compared these odds ratios to those of the three most common oncogenic alleles of *KRAS* in CRC (i.e. G12D, G12V, and G13D). K-Ras^G13D^ was included because, like K-Ras^A59T/E^, it also activates K-RAS through enhanced nucleotide exchange and has a reduced affinity for RAF (21). In all cases, the Ala59 mutants exhibited statistically significant co-occurrence with NF1 and member of the RTK family, while tumors harboring KRAS G12D, G12V, and G13D mutations were more likely to not have a secondary mutation (**Fig. 3C**). Consistent with previous studies, cancers expressing K-RAS^G13D^ did favor loss of NF1, but surprisingly tumors expressing K-RAS^G12D^ did not exhibit mutual exclusivity with EGFR, despite previous work showing strong negative selection between K-Ras^G12D^ and EGFR activation in mouse models of lung cancer (11,26). Altogether, these data demonstrate that K-RAS^A59T/E^, likely because of their inability to robustly activate B-RAF, require cooperating mutations that enhance their ability to activate MAPK signaling.

### NF1 loss cooperates with K-Ras^A59E^ expression in vivo

We discovered that NF1 loss-of-function mutations/deletions commonly co-occur in cancers harboring either K-Ras^A59T^ or K-Ras^A59E^ (**Fig. 3B**). To test if K-Ras^A59E^ cooperates with loss of Nf1, we combine our colon homeostasis and tumor models with a Cre-dependent conditional allele of Nf1 (27). The effect of Nf1 loss in the mouse colon and in models of colon cancer has not previously been reported. We found that loss of Nf1 in the mouse colon promotes weak hyperplasia (**Supplementary Fig. S3A**) and, in the context of colon tumors lacking Apc, both complete and heterozygous knockout of Nf1 significantly decreased survival compared to mice that retain both copies of functional Nf1 (**Supplementary Fig. S3B**). These data provide experimental support for a role of Nf1 in suppressing proliferation in colonic epithelial cells and in colorectal cancers.

When combined with K-Ras^A59E^, knockout of Nf1 in the colonic epithelium resulted in a hyperplastic phenotype statistically indistinguishable from Nf1 null (**Fig. 4A**). When combined, K-Ras^A59E^ and Nf1 loss cooperate to produce an extreme hyperplastic phenotype similar to expression of K-Ras^G12D^. Similarly, in the colon tumor model, Nf1 loss cooperated with K-Ras^A59E^ to reduce mouse survival to a similar extent as K-Ras^G12D^ (**Fig. 4B**). However, combined mutation of K-Ras and Nf1 did not increase colon dysplasia and growth relative to mice that only expressed K-Ras^A59E^ (**Fig. 4C,D**). Likewise, we did not see a statistically significant increase in MEK or ERK phosphorylation when Nf1 was mutated in combination with K-Ras^A59E^ (**Fig. 4E,F**). However, we did note that knockout of Nf1 in the colon resulted in an increase in AKT phosphorylation, but this molecular phenotype was lost in Apc-mutant tumors (**Supplementary Fig. S3C**).

**Figure 4.**
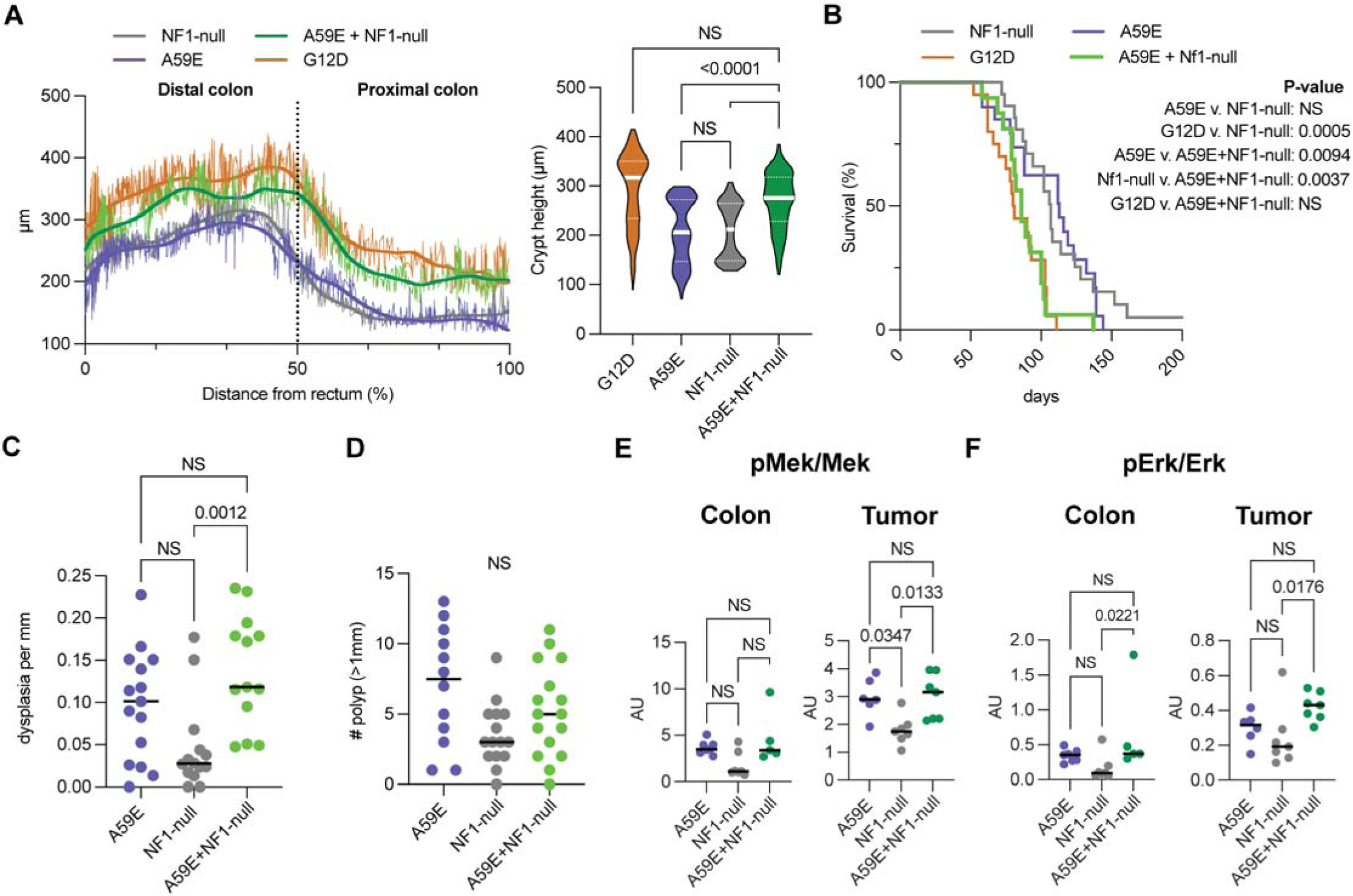
KRAS A59E is further activated in NF1-null tissues. (A) Crypt heights by distance from the anus to the cecum. Left, thick line represents the calculated LOWESS line using a 10-point smoothing option. The thin jagged line represents a smoothened 2nd order polynomial calculated at every 10 points. Right, quantification of crypt lengths. (B) Survival curve of Fabp-cre(+), Apc(+/2lox14) mice either wildtype for K-Ras (N=60), or harboring LSL-K-Ras G12D (N=19), LSL-K-Ras A59E (N=20) alleles, Nf1(fl/fl) (N=19), or both LSL-KRas A59E and Nf1(fl/fl) together (N=17). P-values are from Log-Rank Mantel Cox tests for survival. (C) Quantitation of dysplasia in colon sections from different tumor mice. (D) Polyps greater than 1mm observed in tumor mice. (E-F) Quantitative western blot analysis of (E) MEK phosphorylation and (F) ERK phosphorylation from mouse colon scrapes and tumors. Statistical analysis of panels (C-F) was performed using Kruskal-Wallis tests with significance cutoff at <0.05.

To explore the role of MEK and ERK signaling in more detail, we assayed Paneth cell formation in the ilium of our mice. K-Ras^G12D^ inhibits Paneth cell differentiation in a MEK dependent manner (12,28). In contrast, K-Ras^A146T^, which weakly activates MEK, does not inhibit Paneth cell differentiation (12). Similar to K-Ras^A146T^, the ilium of K-Ras^A59E^ mice and Nf1 knockout mice exhibited normal Paneth cell formation (**Supplementary Fig. S3D**). Since MEK and ERK phosphorylation are unchanged in tissues with K-Ras and Nf1 mutations, we also anticipated that these mice would develop Paneth cells. However, expression of K-Ras^A59E^ in the context of Nf1 loss inhibited Paneth Cell development (**Supplementary Fig. S3D**). Thus, our data argue against a simple quantitative model of MEK/ERK activation, but rather the activation of distinct RAF/MEK/ERK axes, as regulating oncogenic-like phenotypes.

### Nf1 regulates the active state of KRAS A59E

To explore the mechanism of cooperation between K-Ras Ala59 mutants and Nf1 loss, we began by measuring K-Ras activation in colons of differing genotypes. Consistent with the weak cellular and tissue-level phenotypes, loss of Nf1 did not increase Ras activation *in vivo* (**Fig. 5A**, **Supplementary Fig. S3E**). Also consistent with our prior biochemical work, K-Ras^A59E^ induced only a small increase in Ras-GTP in mouse colons (**Fig. 5A**). We did, however, observe a specific increase in K-Ras^A59E^ activation in both colonic epithelium and tumors in response to Nf1 knockout. That the increased Ras-GTP is K-Ras^A59E^, and not WT Ras, is apparent from the slow electrophoretic mobility of the mutant protein. By extension, loss of Nf1 specifically activates K-Ras^A59E^ protein.

**Figure 5.**
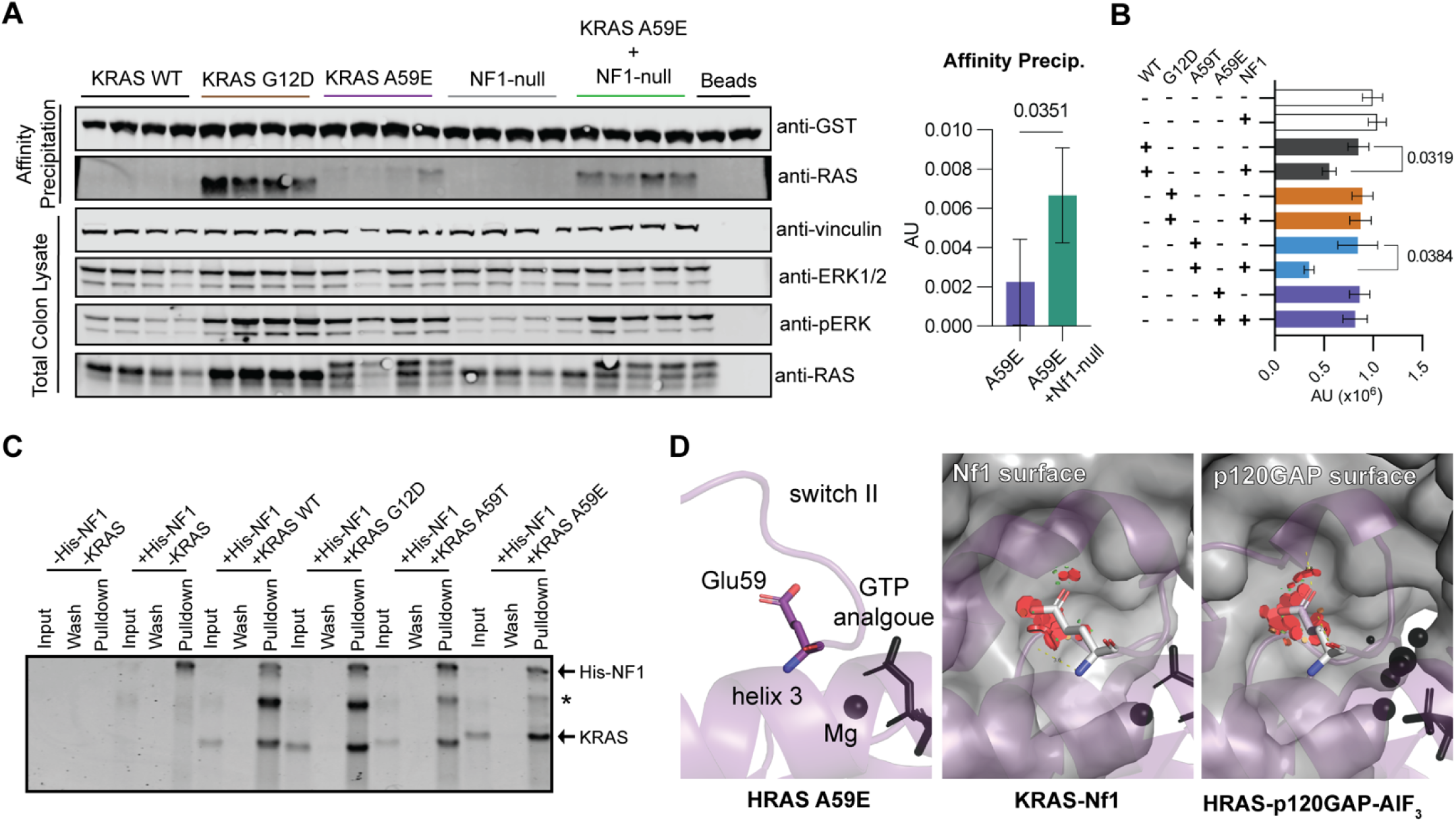
NF1 binds to KRAS A59E. (A) Left, western blot results from Ras activity assay using GST-CRAF-RBD. Right, quantitation of KRAS A59E pulldown by GST-CRAF-RBD from colon scrapes of Fabp-Cre(+), K-Ras(+/LSL-A59E) and Fabp-Cre(+), K-Ras(+/LSL-A59E), Nf1(fl/fl) mice. P-values are the result of unpaired t-test between groups. (B) Results of GTP hydrolysis by NF1 and labelled KRAS proteins. Bars represent the average of three experiments each with 5 technical replicates. Error bars are standard deviation and statistics are the result of a paired t-test. Note that wildtype KRAS without NF1 showed significant intrinsic hydrolysis compared to both non-RAS +/− NF1 control wells (p-values 0.0012 and 0.0138, respectively). (C) Pulldown of purified wildtype KRAS4B bound to GppNHp and its mutants G12D, A59T, and A59E by the His-tagged catalytic domain of NF1. * Indicates remaining uncleaved 6xHis-KRAS4B. A repeat of this experiment is shown in Supplemental Figure 3E. (D) Modeling of A59E mutation effects in RAS/GAP interactions. Left, crystal structure of HRAS A59E (PDB code 7JIH) bound to non-hydrolysable analogue of GTP and preferred rotamer of Glu59 when RAS A59E is in the active state. Middle, crystal structure of KRAS bound to a non-hydrolysable analogue of GTP and in complex with the catalytic domain of NF1 (PDB code 6V65). Right, crystal structure of HRAS bound to GDP and a transition state mimic of GTP hydrolysis in complex with the catalytic domain of p120GAP (PDB code 1WQ1). Nucleotide analogues, transition state complex, and active site magnesium ions are colored black and the surfaces of the NF1 and p120GAP catalytic domains are shown as gray surfaces. Modelled Glu59 is shown in light grey in the middle and right panels. Red disks represent clashes calculated by PyMol during *in silico* mutagenesis.

We previously demonstrated that K-Ras^A59E^ was insensitive to the GAP catalytic domain of RASA1 (*i.e.* p120GAP) (13). However, the mechanisms by which RASA1 and NF1 catalyze GTP hydrolysis are distinct (11,29) and we found that purified K-Ras^A59T^, but not K-Ras^A59E^, was sensitive to Nf1-mediated GTP hydrolysis (**Fig. 5B**) despite both mutants retaining the ability to associate with the NF1 catalytic domain (**Fig. 5C**, **Supplementary Fig. S3F**). Our data are consistent with previous experiments showing that Nf1 and K-Ras^A59E^ can associate in the cell (2). Modeling of Glu59 in the active site of KRAS bound to the catalytic domain of Nf1 using *in silico* mutagenesis and a rotamer of glutamate similar to its conformation found in our crystal structure of H-Ras A59E bound to GppNHp support that, while Glu59 will undoubtedly reduce the affinity between K-Ras and Nf1, it can be accommodated in a pocket in the final K-Ras-Nf1 complex with minimal clashes (**Fig. 5D, Left versus middle panel**). While this pocket was also present for the H-RAS-p120GAP complex, more extensive clashes are made between p120GAP and Glu59 (**Fig. 5D, Left versus right panel**).

GAPs compete for Ras binding with effectors, such as RAF kinases (30,31). Our data suggest a unique model of K-Ras activation by A59E mutation and co-inactivation of Nf1, whereby Nf1 inactivation facilitates more ‘freely’ activated K-Ras to interact with downstream effector proteins (**Fig. 5E**). Note that this model is distinct from KRAS the action of G12D mutation, which unlike A59E promotes nucleotide exchange in the direction of the active GTP bound state, whereas A59E does not (**Fig. 5E**) (13). Thus, it is possible that NF1 antagonizes KRAS A59E association with ARAF, and with its deletion ARAF stabilizes the GTP bound form of KRAS A59E (32). Regardless, our data show that NF1 knockout directly synergizes with KRAS A59E to promote more severe cancer-like phenotypes of the mouse colon.

### K-Ras^G12D^ and K-Ras^A59E^; Nf1-null genotypes show overlapping transcriptomes

To better understand the biology driving differences in autochthonous tumor growth in our different genotypes, we performed bulk RNA-seq. After merging of datasets (**Supplementary Fig. S4A**), principal component analysis exhibited defined separation of K-Ras wildtype and K-Ras^G12D^ tumors, while K-Ras^A59E^, Nf1-null, and the doubly mutant mice were intermediate along the first principal component (**Supplementary Fig. S4A**). However, tumors expressing K-Ras^A59E^ in the background of Nf1 knockout exhibited the largest degree of variation along the second principal component axis (**Supplementary Fig. S4A**). Considered in the context of the phenotypic data in **Fig. 4**, the gene expression data indicate that K-Ras^G12D^ and K-Ras^A59E^; Nf1 null colon are phenotypically similar, by quite different at the molecular level. Indeed, comparison of differentially expressed genes in the five genotypic groups indicates that all tumor types diverges from the transcriptome of K-Ras^G12D^ colon tumors (**Supplementary Fig. S4B**). However, we did note that variation in gene expression between doubly mutant and K-Ras^G12D^ tumors was less than that of K-Ras^A59E^ or NF1-null tumors (**Supplementary Fig. S4B,C**).

To identify shared biological processes that enhance tumor growth, we performed Gene Set Enrichment Analysis (GSEA) using transcriptomes from our different tumor groups. Compared to tumors expressing WT K-Ras, K-Ras^G12D^ tumors exhibited dysregulated transcriptomic hallmarks of cell growth, survival, and immunity, including signatures related to K-Ras signaling (**Supplementary Fig. S4D**). In contrast, GSEA analysis comparing tumors with WT KRAS against those harboring K-Ras^A59E^ or lacking NF1, or their combination, demonstrated that the doubly mutant genotype shared only one dysregulated hallmark with K-Ras^G12D^ that was not found in other genotypes: p53 pathway regulation (**Supplementary Fig. S4D**). Upregulation of p53 related signaling in K-Ras^G12D^ and doubly mutant tumors makes sense, as these two genotypes would be expected to induce cellular stress (33,34). We also noted that K-Ras^A59E^ or NF1-null tumors favored enrichment of transcripts related to E2F function and cell growth, but doubly mutant tumors showed de-enrichment of E2F targets of transcription (**Supplementary Fig. S4D**). Thus, while we expected an additive effect on the tumor transcriptome, we instead observed that tumors expressing K-Ras^A59E^ in the context of Nf1 loss are as different from K-Ras^G12D^ as they are from K-Ras^A59E^ and Nf1-null tumors.

Since the transcriptomic signatures of doubly mutant tumors and K-Ras^G12D^ do not converge, but their DEG’s do (**Supplementary Fig. S4B,C**), we reason that these DEGs reflect a minimum necessary change in the transcriptome necessary to induce the oncogenic phenotypes seen in our mice (**Fig. 4**). To identify genes, rather than gene sets, that were regulated similarly in doubly mutant tumors and tumors expressing K-Ras^G12D^, we performed an intersection analysis. We identified ~60 genes from doubly mutant tumors that were dysregulated in a similar manner as K-Ras^G12D^ (**Supplementary Table S2**) and further interrogation of these genes using the Dependency Map (35) identified the molecular chaperone HSPA8 (also known as HSC70) as important for K-Ras mutant CRC cell growth, but not CRC cells expressing WT KRAS (**Supplementary Fig. S4E**). While merging of our bulk RNA-seq datasets reduces the sensitivity of our analysis, enhanced expression of HSPA8 makes sense as this chaperone is known to stabilize CCND1 and is a component of the functional CCND1/CDK4 complex (36). Thus, combined signaling through KRAS caused by G12D mutation alone, or by combined mutation of A59E in combination with NF1 loss, may contribute to cell cycle progression through CCND1 stabilization at multiple levels (37), primarily through ARAF and CRAF.

### K-Ras^A59E^ remains dependent on upstream and downstream EGF receptor signaling

In contrast to Gly12 mutants of K-Ras, observations from the clinic suggest that patients whose cancer expressed K-Ras^A59T/E^ may benefit from therapies targeting EGFR (4,38). To test this hypothesis, we generated organoids from the colon epithelium of Fabp-Cre(+) mice expressing WT, G12D, or A59E K-Ras. Since cetuximab and panitumumab recognize human EGFR, we used a ‘murinized’ version of cetuximab (mCetux) that recognizes mouse EGFR to experimentally determine whether cancers expressing K-Ras^A59E^ are indeed sensitive to EGFRi (39). Consistent with clinical observations, organoids expressing K-Ras^A59E^ are sensitive to mCetux relative to those expressing K-Ras^G12D^ (**Fig. 6A**). We also used a small molecule kinase inhibitor, Erlotinib, as well as SHP2 (i.e. RMC4550), and MEK1/2 (i.e. Trametinib) inhibitors to test the relationship between EGFR signaling and K-Ras^A59E^. Consistent with a dependence on upstream EGFR function and our mCetux experiments, Erlotinib inhibited K-Ras^A59E^ organoid growth and provoked cytotoxicity, in contrast to colon organoids expressing K-Ras^G12D^ (**Fig. 6B**). Likewise, K-Ras^A59E^ organoids were modestly sensitive to the SHP2 inhibitor RMC-4550 (**Fig. 6C**), while not differing from other genotypes in response to MEK inhibition (**Fig. 6D**). These data imply that K-Ras^A59E^ is still generally sensitive to EGFR activation. By extension, we measured the ability of colonic organoids to respond to extracellular EGF and found that, unlike K-Ras^G12D^ organoids, K-Ras^A59E^ organoids proliferate in response to extracellular EGF (**Fig. 6E**). Altogether, these data show that cancers expressing K-Ras^A59E^ organoids are retain dependence on upstream EGFR signaling.

**Figure 6.**
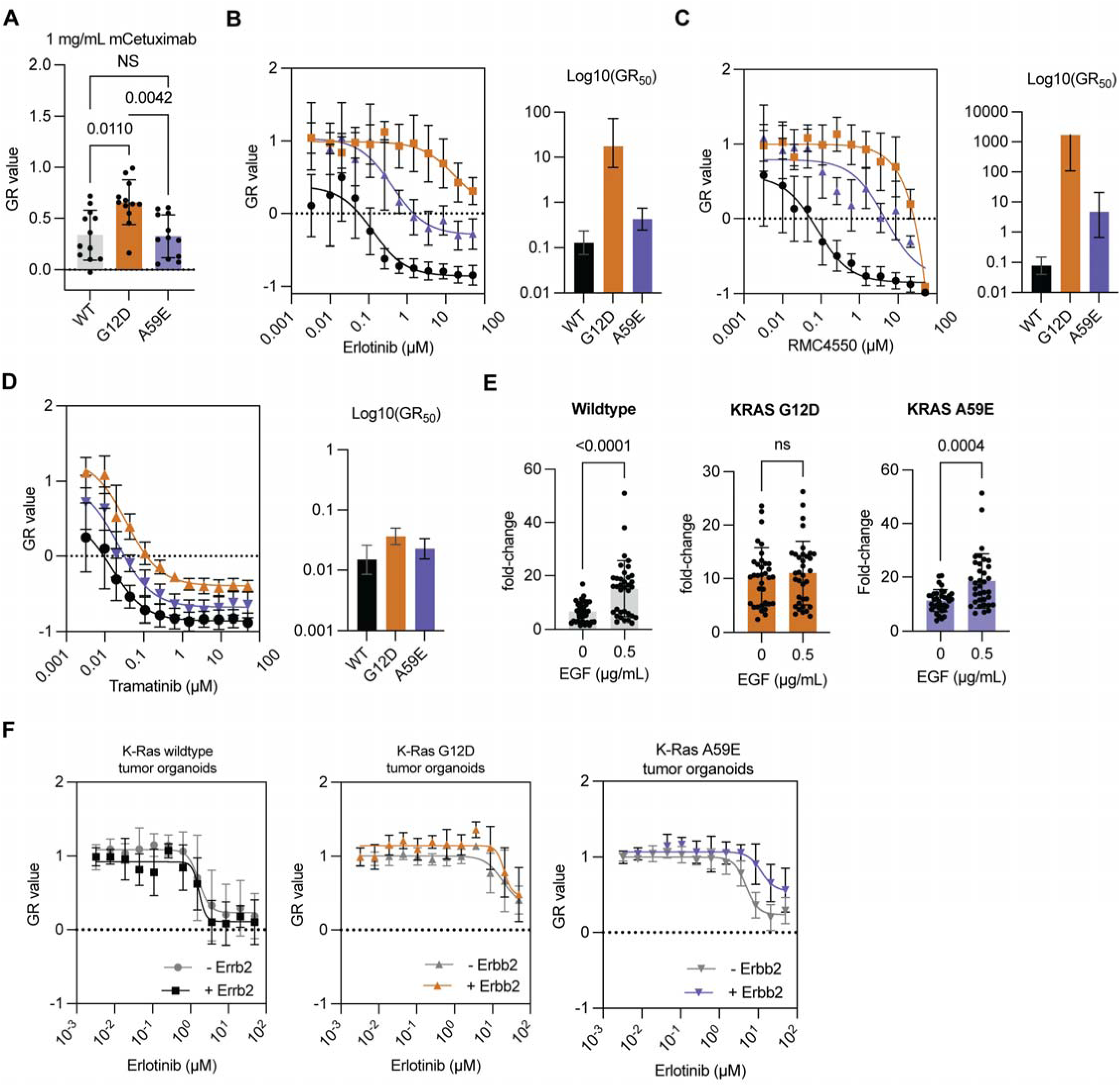
KRAS A59E organoids are sensitive to upstream inhibition of MAPK signaling. (A) GR values for colon tumor organoids at 1mg/mL of mCetuximab. (B-D) Growth normalized (GR) values for colon organoids exposed to different concentrations of Erlotinib (B), RMC4550 (C), and Tramatinib (D) over 7-days. Left panels, drug titration curve. Right panels, calculated GR50 value from curve on the left expressing different KRAS mutants. Colors on left panels are matched on right panels. (E) Growth of colon organoids after 7 days with and without EGF in media. ‘Fold-change’ refers to increase in PrestoBlue metabolism from day 7 compared to day 1. (F) Colon tumor organoids expressing KRAS G12D, KRAS A59E or wildtype KRAS infected with lentivirus to express ERBB2 or empty vector exposed to different concentrations of Erlotinib over 7-days. Statistics are generated via a Kruskal-Wallis tests.

Our prior bioinformatic analysis identified other EGFR and other RTKs as commonly co-mutated with K-Ras^A59T/E^ in primary colorectal cancer (**Fig. 3B**). A consequence of these co-mutations could be resistance to EGFRi (40). To test this possibility, we examined whether ERBB2 over-expression (*i.e.* amplification) conferred resistance of EGFRi upon organoids expressing K-Ras^A59E^ (**Supplementary Fig. S5B-D**). Expression of ERBB2 in tumor organoids expressing K-Ras^A59E^ promoted resistance to Erlotinib, whereas expression of ERBB2 did not appear to alter the response of K-Ras WT or G12D tumor organoids (**Fig. 6F**). Since the level of ectopic expression of ERBB2 in our tumor organoids was not sufficient to promote resistance to Erlotinib in organoids WT for KRAS, we reason that resistance in the K-Ras^A59E^ tumor organoids must be the result of cooperation between K-Ras^A59E^ and ERBB2. Thus, our data establish that cancers expressing K-Ras^A59T/E^ are likely sensitive to EGFRi, unless that carry a cooperating gene in an RTK.

## Discussion

Like other strongly activating mutants of K-Ras, A59T and A59E mutations suppress intrinsic GTP hydrolysis and enhance nucleotide exchange (13). Thus, one would expect that these K-Ras mutants would be strongly oncogenic, but that is not the case. Unlike other K-Ras mutants, A59T and A59E paradoxically inhibit RAF localization, association, and activation, and we show here that these effects are most significant for B-RAF (13,16,17). But, despite their attenuated capacity to promote RAF activation, we show using genetically engineered mice that K-Ras^A59E^ is a bo*na fide* oncogenic allele, as it promotes hyperplasia in the mouse colon and cooperates with Apc loss to reduce mouse survival. Thus, A-RAF and C-RAF appear limited in their ability to drive hyperplastic and oncogenic phenotypes, while B-RAF dimerization and activation appears to play a unique role in colorectal tumorigenesis. Our conclusions are supported by tumor mutation patterns. Of the top three cancers that favor K-Ras mutation (*i.e.* PDAC, NSCLC, and CRC), mutations in B-RAF are 2-5 times more likely in CRC (11% vs. 6% in NSCLC and 2% in PAAD), and B-RAF mutations in CRC favor activating V600 mutations rather than non-V600 and cooperative mutations (68% in CRC vs. 26% in NSCLC and 22% in PAAD) (41–43). Thus, we unintentionally demonstrate that the K-Ras^A59E^ mouse model is an *in vivo* system by which RAF dimerization, in the context of K-Ras function, can be interrogated in better detail. This contrasts with typical K-Ras hyperexchange mutants (*i.e.* G13D and A146T), which have been studied in mice and which promote mild colonic phenotypes like K-Ras^A59E^, but which do not show defects in B-RAF dimerization (9,12,21,44).

Our data support a strong role for mutant-specific K-Ras functions in tumor evolution (23), as a deficiency in RAF signaling provides a causal role for the unique range of MAPK activating co-mutations seen in cancer expressing K-Ras^A59T/E^. In support of this concept, we observed that expression of K-Ras^A59E^ in the context of Nf1 loss produces a collection of more severe phenotypes. A necessity for co-mutations might also explain why K-Ras^A59T/E^ are favored in cancers of the gastrointestinal tract, which frequently show high mutation rates, high *KRAS* allele heterogeneity, and long life-histories (1,7). Notably, neither K-Ras^G13D^ nor K-Ras^A146T^ favor the same rates of co-mutation in genes that regulate MAPK signaling as do K-Ras^A59T/E^. However, while co-mutation of Nf1 knockout increased the oncogenicity of K-Ras^A59E^, these mice did not completely phenocopy K-Ras^G12D^ at the molecular level. Instead, differences in cellular signaling between doubly mutant mice and K-Ras^G12D^ resulted in distinct transcriptomes, underscoring the unique and significant role that K-Ras plays controlling cell growth. However, our experiments did identify a potentially shared feature between doubly mutant mice and K-Ras^G12D^: upregulation of HSPA8 gene expression. HSPA8 has a particular role in the stabilization of cyclin D1 to support of cell cycle progression, as well as broader functions as a cellular chaperone for protein folding. Thus, strong oncogenic signaling from K-Ras may depend on upregulation of HSPA8 expression and stabilization of CCND1. However, support for HSPA8 as a mediator of oncogenic KRAS function remains to be tested *in vivo*.

Cooperation between Ala59 alleles of K-Ras and other genes of the MAPK signaling pathway in cancers has been observed previously (11,24,43,45–47), but, perhaps, the oldest analogous phenomenon is seen in murine tumor-promoting retroviruses (13). A59T allows Ras proteins to undergo autophosphorylation (p-K-Ras^A59T^) (13). Viral homologues of K-Ras and H-Ras differ from cellular Ras in that Gly12 and Ala59 are substituted, with both carrying A59T. C*is* co-mutations in K-Ras (*e.g.* G12V/A59T) activate the non-phosphorylated form of K-Ras^A59T^ while increasing the steady-state concentration of p-K-Ras^A59T^, and K-Ras^G12V/A59T^ promotes greater cell transformation than either mutant alone (13). Consistently, K-Ras co-mutations are the third most common form of co-mutation in tumors expressing K-Ras^A59T^ (**Fig. 3B**). Thus, inactivation of NF1 can be seen as an alternative to co-occurring Gly12 mutations, as its knockout would be expected to increase the active form of K-Ras^A59T^ (**Fig. 5B**).

The *in vivo* synergy between Nf1 knockout and K-Ras^A59E^ could occur through various mechanisms. An attractive speculation is that NF1 knockout rescues BRAF dimerization, via activation of wildtype HRAS and NRAS, to complement the attenuated RAF signaling promoted by KRAS A59E. Biochemically, NF1 does not have preference for the different isoforms of RAS (48). In fact, the increase in AKT phosphorylation we observed in the mouse colon epithelium could be the result of H- and NRAS activation (49). However, differences AKT phosphorylation are lost in tumors, and we see the most significant changes in RAS isoform activation at the level of mutant KRAS but not wildtype RAS. Furthermore, a recent study showed that NF1 loss does not induce a significant increase in BRAF-related dimers in melanoma cells, arguing against the rescue of BRAF dimerization via pan-RAS activation (50). Alternatively, NF1 has been shown to regulate the GTP bound state of MRAS (51). Thus, loss of NF1 could upregulate MRAS-GTP to stabilize the MRAS-SHOC2-PP1C complex, which regulates CRAF activity via removal of an inhibitory phosphorylation (6,52). Thus, NF1 knockout may potentiate RAF kinase activation through other GTPases. This scenario is consistent with our western blot and transcriptomic studies showing that tumors from doubly mutant mice and KRAS^G12D^ do not strongly converge at the molecular level. However, a more recent report on MRAS biochemistry presents a conflicting view of MRAS, whereby this GTPase is trapped in its GDP bound state, which would likely preclude an NF1/MRAS regulatory axis (53). A third mechanism, and one that is better supported by our data, is that K-Ras^A59E^ is directly activated, as measured by GTP binding state, by Nf1 knockout. This was a surprising result as the catalytic domain of NF1 does not promote GTP hydrolysis on K-Ras^A59E^, although it does for K-Ras^A59T^ (**Fig. 5B**). Instead, NF1 may act as an antagonist of ARAF-KRAS association, as suggested by recent studies from the Rosen lab (32). In this model, loss of NF1 facilitates an increase in the steady-state binding of ARAF to KRAS^A59E^. Mechanistically, the ARAF/KRAS^A59E^ complex, which is more stable than the NF1/KRAS^A59E^ complex, prevents nucleotide equilibration of KRAS^A59E^ and could enhance the concentration of GTP bound KRAS^A59E^ in the cell (**Figure 7**). Interestingly, from our data we expect KRAS^A59T^ to behave somewhat differently. Unlike KRAS^A59E^, KRAS^A59T^ remains sensitive to NF1. Thus, KRAS A59T is likely activated by both a reduction in GTP hydrolysis and by potential titration of the activated state of its phosphorylated proteoform. Regardless, our data suggest that our understanding of how the cell regulates the activated state of KRAS is incomplete, with other unknown mechanisms playing a role in the activation of KRAS.

**Figure 7.**
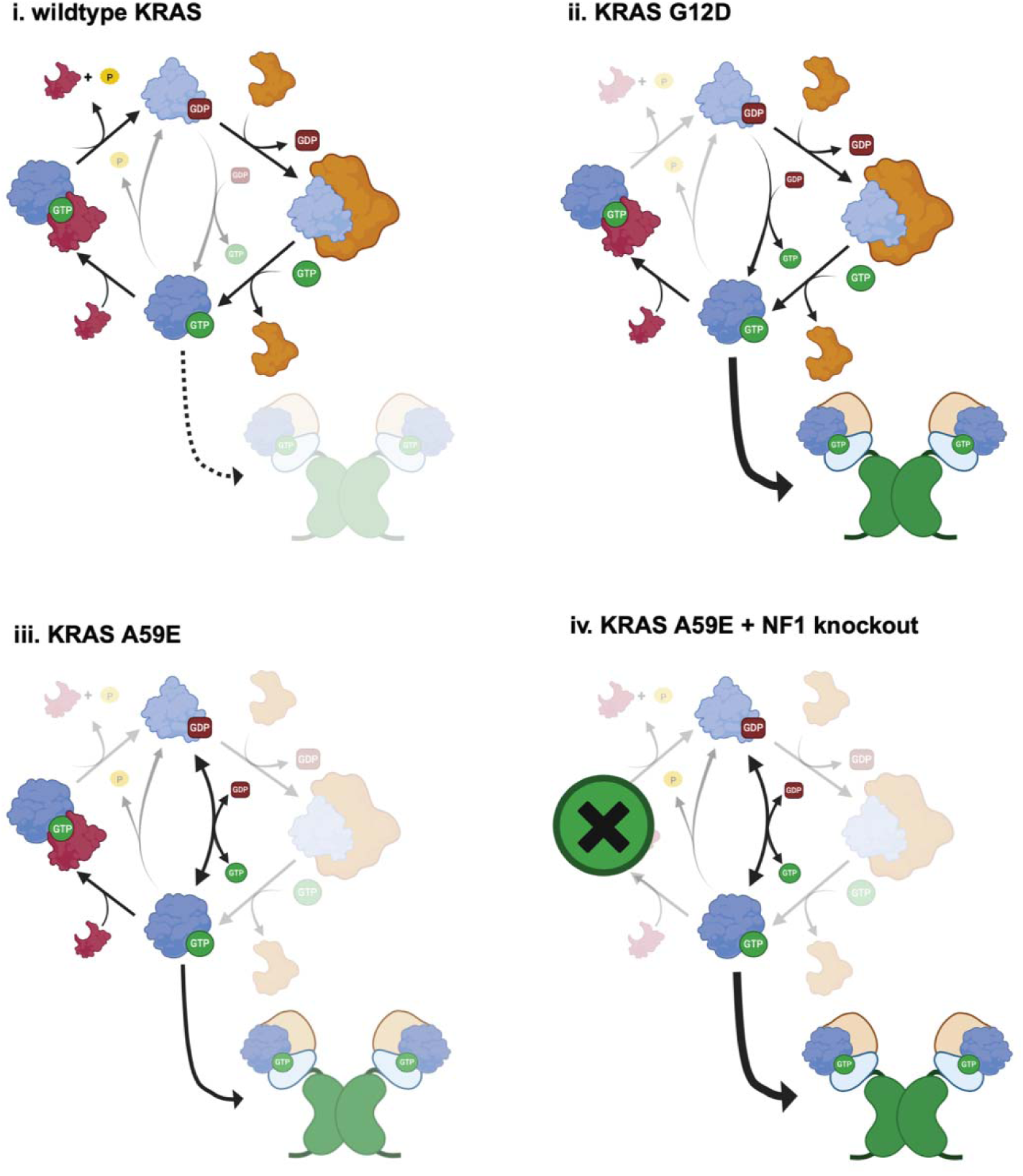
Model of KRAS^A59E^ activation by NF1 inactivation. Model of GTPase cycle for (i) wildtype KRAS, (ii) KRAS G12D, (iii) KRAS A59E, and (iv) KRAS A59E in the context of NF1 knockout. Note that the intrinsic pathways of nucleotide exchange (left side of each panel) is different for the G12D and A59E mutants, and are based on data from (13). Dimeric structure opacity refers to activation of RAF kinases through KRAS binding to RBD and CRD domains. Figure was created with BioRender.com.

Altogether, our data suggest that tumors harboring K-Ras^A59T/E^ mutations, along with their secondary mutations, represent a unique sub-type of K-Ras-mutant colorectal cancer. Unlike other oncogenic K-RAS mutations, tumors harboring A59T and A59E alleles of K-RAS do not show strong intrinsic resistant to EGFRi, and we provide experimental support for clinical observations (3,4). Study of EGFRi sensitivity in both tumor and colon organoid culture, in the context of K-Ras^A59E^ signaling indicate that B-RAF function may be important for oncogenic K-Ras to provide EGFR independence and intrinsic resistance to EGFRi in CRC cells. Consistently, overexpression of ERBB2, which promotes B-RAF/C-RAF dimerization (54), in our colon tumor organoid experiments promoted resistance to EGFRi in the context of K-Ras^A59E^. Class 3 B-RAF mutations, which amplify upstream signaling in the absence of co-occurring resistance alterations, also show a limited capacity to promote intrinsic resistance to EGFRi due to their inability to promote B-RAF dimerization without the aid of K-Ras mutation (43,45,46). Thus, CRC with K-Ras A59 mutations are perhaps similar to class 3 B-RAF mutations in that they also could respond to EGFR inhibition if they lack a co-occurring second mutation in the RTK/RAS pathway. Thus, our data argue for a broader experimental assessment of EGFRi resistance for oncogenic K-Ras mutant, alone or in combination with co-occurring RTK/RAS pathway mutations, in order to better capitalize on the use of EGFRi in colon cancers.

## Methods

### Non-embryonic stem cell tissue culture

All mammalian cells (*except mES cells, see below*) were maintained at 37°C in a humidified incubator with 5% CO_2_ and 95% humidity. HeLa cells were cultured in EMEM containing 1% Pen/Strep, 10% Fetal Bovine Serum (FBS). HEK293T cells were cultured in DMEM supplemented with 1% Pen/Strep, 10% FBS. HEK293T and L-WRN cells were purchased from ATCC. Mammalian cells in two-dimensional culture were maintained using a 3t3 culture protocol. L-WRN cells were propagated and used to make conditioned medium as previously described (55).

Description of mouse colon and tumor organoid derivation is described in the methods below. Mouse colon organoids were cultured in a Matrigel matrix of 10,000 cells per 50µL dome (Thermo Fisher Scientific) and submerged in 500µL of colon organoid media containing Advanced D-MEM/F-12 culture medium, L-WRN-conditioned medium, 1% (v/v) B-27 supplement (Gibco), 1% (v/v) GlutaMax (Gibco), 10mM HEPES (Gibco), 10mM nicotinamide (Sigma), 0.5% (v/v) N-2 supplement (Gibco), 500µM N-Acetylcysteine (Sigma), 50nM [Leu15]-Gastrin I (Sigma), 500nM A83-01 (Sigma), 10µM SB202190 (Sigma), 50ng/mL recombinant human EGF (Gibco), 1µM PGE2 (Sigma), and 100µg/mL Primocin (InvivoGen). Organoids were passaged once a week by adding 55µl of Dispase (Corning) per well directly followed by a ~2 hours digestion at 37°C. Two rounds of TrypLE (Gibco) digestion was used to generate a single cell suspension of colon organoids. Mouse colonic tumor organoids were cultured as above, except they were maintained in the Wnt depleted media composed of Advanced D-MEM/F-12 medium, 1% (v/v) B-27 supplement, 1% (v/v) GlutaMax, 10mM HEPES, 0.5% (v/v) N-2 supplement, 500µM N-Acetylcysteine, 50ng/mL recombinant human EGF, 100ng/ml mouse Noggin (PepProtech) and 100µg/mL Primocin.

### Co-immunoprecipitation experiments of RAF isoforms

Experiments were performed as previously (13,44). HeLa cells were plated at a concentration of 8.3×10^5^ per 10 cm dish. 18-24 hour prior to transfection, pCMV5-Venus-K-RAS constructs were transfected into cells using the XtremeGENE9 transfection reagent kit at a 2:1 ratio of XtremeGENE9 to DNA. 48h post transfection, serum-starved cells were washed twice with ice cold PBS and lysed in1% NP-40 lysis buffer (20mM Tris [pH 8.0], 137 mM NaCl, 10% glycerol, 1% NP-40 alternative, 0.15 U/mL aprotinin, 1 mM PMSF, 0.5 mM sodium vanadate, 20 mM leupeptin; 500mL/10 cm dish) for 15 min at 4°C. Lysates were clarified by centrifugation at 14,000 RPM for 10 min at 4°C. Protein content was determined by Bradford assay. Lysates containing equivalent amounts of protein were incubated with either anti-C-RAF (BD Pharmagen, 610152) or anti-A-RAF (Santa Cruz, sc-408) and protein G Sepharose beads for 2h at 4°C on a rocking platform. Complexes were washed with 1% NP-40 buffer and examined by western blot analysis in tandem with equalized total cell lysate. The following antibodies were used for blotting: ARAF (Santa Cruz, sc-408), BRAF (Santa Cruz, sc-5284), CRAF (BD Pharmagen, 610152), GFP (MBL International, D153-3), Venus (Roche, 118114460001), MEK1 (BD Biosciences, 610122), and phospho-MEK1/2 (Cell Signaling Technologies, 9121).

### Mining of tumor genome samples and related analyses

Determination of co-mutations in genes that regulate the MAPK signaling pathway in human colorectal tumors already harboring different mutant KRAS alleles was performed using a previously compiled dataset from Cook et al. (23). Genes selected for co-mutation analysis were those defined as the RTK/RAS pathway genes from The Cancer Genome Atlas (TCGA) (22). Methods for this analysis are described in (23). For analysis of enrichment of Ala59 mutation per cancer, the frequency of Ala59 mutations in GI and blood cancers, and identification of co-mutations, a new dataset comprising public tumor samples from cBioPortal and GENIE databases (v15.0-public) was created (41,42,56). Tumor samples from non-public sources (Memorial Sloan Kettering, MD Anderson) also contributed to this dataset (**see Supplementary Table 1**). Data presented in **Figure 3** is for putative driver mutations in the RTK-RAS pathway only. For the log odds ratio analysis, data were taken exclusively from public colorectal tumor samples available through GENIE (v15.0-public). To reflect genetic interactions between KRAS alleles and RTK family members (EGFR, ERBB2, ERBB3, and ERBB4), NF1, NRAS, and BRAF, calculated odds ratios reflect differences between tumor samples, rather than patient samples (i.e. in some cases multiple tumor samples are present for a single patient). While assignment of KRAS mutation was allele specific (i.e. G12D, G12V, G13D, A59T, A59E), RTK genes, NF1, NRAS, and BRAF were assessed for somatic mutations, structural variants, and copy-number alterations. Generation of chi-square tables was done in excel. Calculation of the odds ratio and statistical significance were done in PRISM (v10) using the Chi-square Yates’ correction test. 95% confidence intervals were also calculated in PRISM using the Gart adjusted logit interval. Visualization of all data was done using PRISM (v10).

### SaCas9 CRISPR mutagenesis of 129/Sv mouse embryonic stems cells

J1 embryonic stem (mES) cells (129S4/SvJae genetic background) containing a conditional endogenous *Kras* allele in which a transcriptional stop element (lox-stop-lox, LSL) cassette between *Kras* exons 1 and 2 were altered using SaCas9 CRISPR mutagenesis to conditionally express K-Ras^A59E^ mutant. All mES cells were culture on the layer of feeders either generated in house or purchased from Applied StemCell with Knockout DMEM media supplemented with 15% FBS (heat inactivated ES cell screened; Hyclone), 1X Glutamax (Gibco), 100µM 2-Mercaptoethanol (Sigma), 1X non-essential amino acid (Gibco), 1X PenStrep, and 10^3^units/mL Leukemia inhibitory factor (LIF; Millipore). mES colonies were grown until visible to the naked eye before splitting. Cell media was refreshed every day, and always 2-3 hours before sub-culture, clone picking, or nucleofection.

Modification of *Kras^LSL-wildtype/+^* mES cells to *Kras^LSL-A59E/+^* was done as described previously (13). Briefly, The SaCas9 plasmid was modified to contain gRNA sequence targeting exon 2 and codon 59 according to the protocol shown in (57). The forward and reverse primers for cloning into the SaCas9 plasmid were 5’-CACCGTTCTCGACACAGCAGGTCAAG-3’ and 5’-AAACCTTGACCTGCTGTGTCGAGAAC-3’. Before nucleofection, mES cells and feeders were washed twice with PBS, and then dissociated by incubation for 10-15 min at 37°C with 0.05% Trypsin-EDTA. Cell solution was collected, excess media was added to inactivate trypsin, and then cells were spun down for 5 min at 4°C at 80xg. Cells were resuspended in media and then applied to a gelatin coated 10 cm dish at 37°C. After 30 min, the non-adherent mES cells were collected and spun down for 5 min at 4°C at 80xg. The cell pellet was resuspended in PBS and cell number was determined using a Nexcelom Bioscience Cellometer^®^ AutoT4 cell counter. Using the same centrifuge conditions, a cell pellet of 3.0×10^6^ K-Ras(+/LSL-wildtype) mES cells was collected and then redissolved in 90 µL of supplemented nucleofector solution prepared per the Lonza Mouse ES cell Nucleofector^®^ kit protocol. Next, the cell solution was gently mixed with 10 µL of 1X TAE containing 40 µg of single-stranded DNA repair template, purchased from Integrated DNA Technologies, and 5 µg of modified SaCas9 plasmid that was purified from *E.coli* using the PureLink® HiPure Plasmid Filter Maxiprep kit. The repair template to generate the A59E substitution and silent restriction site was 5’-CTGTCTGTAATAATCCAGACTGTGTTTCTCCCTTCTCAGGACTCCTACAGGAAACAAGTAGTAATTGATGGAGAAACCTGTCTCTTGGATATCCTCGACACAGAAGGTCAAGAAGAGTACAGTGCAATGAGGGACCAGTACATGAGAACTGGGGAGGGCTTTCTTTGTGTATTTGCCATAAATAATACTAAATCATTTGAAGAT-3’. Nucleofection was performed according to the Lonza Mouse ES cell Nucleofector^®^ kit protocol. Briefly, the mES cell and DNA solution was carefully added to the supplied cuvette using a plastic pipette. Care was taken to limit air bubbles. The cuvette was then placed in a Nucleofector^®^ II device and the program A-013 was run. Immediately after nucleofection, 500µL of pre-warmed media was added to the cuvette and gently mixed using the plastic pipette. The entire solution was then added to single well of a gelatin coated 6-well dish with feeders. Cells were fed fresh media every day for 2-3 days. After 2-3 days, cells were collected, diluted 1:1000, and plated on three gelatin coated 10 cm dishes with feeders. The unused cells were spun down at 4°C for 5 min at 80xg, resuspended in chilled mES freezing media (80% media, plus an extra 10% HyColone FBS and 10%DMSO), and then stored for later use at −80°C.

The colonies with well-defined edges were picked in 96-well plates containing 100 µL of trypsin-EDTA in each well. After incubation at 37°C for 5-10 minute, another 100 µL of complete ES cell media was added to each well, cells were resuspended, and then split between two gelatin coated 96-well plates containing feeders. mES clones were then transferred back to 37°C. After 2-4 days, mES clones in one plate were frozen and stored at −80°C by first washing with PBS, dissociating the cells with 50 µL trypsin-EDTA for 10 min, and then mixing dissociated cells with 50 µL of 2X mES freezing media containing 60% ES cell media, 20% FBS and 20% DMSO. The other 96-well plate underwent genomic DNA extraction using the *Quick*-DNA kit from Zymo Research. The Picked ES cell clones were screened by PCR with Ex2 genotyping primers (forward 5’-TGTGACCATTAGCATTGTTTGC-3’, reverse 5’-CTTAAACCCACCTATAATGGTG-3’), followed by EcoRV enzyme digestion. Positive clones were identified by the presence of an undigested PCR product migrating at ~850 bp and a digested product migrating ~700 bp. Positive clones were further validated by sequencing using the forward or reverse Ex2 sequencing primers, and a touchdown protocol using a T_m_ of 56°C. The PCR product was further confirmed by Sanger sequencing using the forward Ex2 sequencing primer. Expression of K-Ras A59E mutant protein in *Kras^LSL-A59E/+^* mES cells was also validated by detection of a higher migration band visible by western blot.

Prior to ES cell microinjection, *Kras^LSL-A59E/+^* ES cells underwent ‘rescue process’ to ensure totipotency using RESGRO^®^ media according to the media manufactures protocol. ‘Rescued’ mES cells were injected into C57BL/6J embryos in the Transgenic Mouse core at Harvard Medical School (https://immunology.hms.harvard.edu/resources/transgenic-mouse-core). Of the injected ~30 mouse embryos, two mice of 30% and 60% chimerism were successfully bred to C57BL/6J mice. The germline transmission of the *Kras^LSL-A59E^* allele was confirmed by genotyping PCR using Ex2 genotyping primer sets (same protocol as the mES cells screening). *Kras^LSL-A59E^* mouse strain was crossed into C57BL/6J (Jackson Laboratories, Strain number: 000664) for at least 10 generations. Experimental data was collected starting at generation N6.

### Housing, breeding, genotyping, and sacrifice of mouse lines

Mice were housed in a barrier facility and fed ad libitum in a temperature-controlled environment with a 12-hour light/dark cycle. All animal studies were approved by the Institutional Care and Use Committee at Beth Israel Deaconess Medical center and Dana Farber Cancer Center. All animal experiments were done with mice that were ≥95% congenic with the C57BL/6 genetic background. To generate mice expressing different K-Ras in the colon, Fapbl4X@-132-Cre mice (hereafter called Fabp-Cre(+) mice) (19) in a C57BL/6 genetic background were backcrossed to C57BL/6 mice either wildtype for KRAS or harboring silent LSL-K-Ras G12D (58) or LSL-K-Ras A59E (this publication) alleles. Weanlings were genotyped via tail DNA for the presence of Fabp-Cre and the LSL cassette. Mice were also further genotyped for the A59E mutation via EcoRV-HF restriction digest. The Nf1flox mouse was generated and characterized previously (59). Heterozygous Nf1flox mice in the C57BL/6 genetic background were obtained from Jackson Laboratories (strain number: 017640). For all experiments, both male and female Fabp-Cre(+) mice were harvested at ≥8 weeks for histology, DNA, and protein based experiments. Note that K-Ras wildtype mice used as negative controls in all experiments were Fabp-Cre(+).

The Apc2lox14 mouse, a stochastic autochthonous colorectal tumor model, was generated and characterized previously (60). In Apc(+/2lox14) mice, Cre expression via Fabp-Cre facilitates constitutive inactivation of one allele of Apc in the colon, and polyp formation follows via recombination and loss of heterozygosity between the wildtype and Cre-edited alleles (60). Apc(+/2lox14), K-Ras(+/LSL-mutant) mice were crossed with Fabp-Cre(+) mice, either with both parents wildtype for Nf1, or Nf1(flox/flox), to generate mice for our tumor and survival experiments. All progeny heterozygous for Apc2lox14 and positive for Fabp-cre, were included in the survival experiments. The humane endpoint for the survival curve was rectal prolapse, along with signs of anemia (e.g pale toes, pale prolapse) or hunching. Experimental mice to examine the role of different mutations on colon homeostasis or cooperation with Apc2lox14 were sacrificed using cervical dislocation and tissue was quickly harvested. Before fixation or freezing, all tissues were flushed and washed using ice-cold PBS.

Genotyping for Fabp-Cre (Primers: F006 5’-TCGTTGACCATTGCTCTCAG-3’; C031 5’-CATCACTCGTTGCATCGACC-3’; Internal-1-F 5’-CAAATGTTGCTTGTCTGGTG-3’; Internal-1-F 5’-GTCAGTCGAGTGCACAGTTT-3’), the LSL cassette (Primers: SD5’ 5’-AGCTAGCCACCATGGCTTGAGTAAGTCTGCA-3’; Forward 5’-CTCTTGCCTACGCCACCAGCTC-3’; Reverse 5’-GTCTTTCCCCAGCAGAGTGC-3’), Nf1flox (Primers: P1 5’-CTTCAGAC TGATTGTTGTACCTGA-3’; P2 5’-CATCTGCTGCTCTTAGAG GAACA-3’; P3 5’-ACCTCTCTAGCCTCAGGAATGA-3’; P4 5’-TGATT CCCACTTTGTGGTTCTAAG-3’), and Apc2lox14 (Primers: Forward 5’-GATGGGTCTGTAGTCTGG-3’, Reverse 5’-GGCTCAGCGTTTTCCTAATG-3’) was done through polymerase chain reaction (PCR) using the same cycling conditions: Step 1: 95°C for 3 minutes. Step 2: 95°C for 30 seconds, 60°C for 30 seconds, 72°C for 1.5 minutes. Repeat step two 35 times. Step 3: final extension of 5 minutes at 72°C. Detection of the A59E mutation in LSL-K-Ras mice was done through PCR and restriction digest. Generation of PCR product for EcoRV-HF digestion was done using the Ex2 genotyping primers and PCR conditions described above. All PCR and digestion products were separated by size using ~1.8% agarose suspended in Tris-acetate-EDTA (TAE) buffer containing SYBR^TM^ Safe DNA stain. Visualization of DNA bands was done using a Gel Doc XR+ from Bio-Rad.

### Colon and tumor tissue preparations for histology, protein, and nucleic acid analysis

Dissection of mice was performed immediately after cervical dislocation. Once euthanized, mice were laid on their back and their torsos were coated with 70% Ethanol. The skin was subsequently raised, a single transverse cut of ~1.2 cm was made, and the skin was pulled in opposite directions along the coronal plane. Next, the peritoneum was carefully cut along the sagittal and transverse planes and the lower gastrointestinal track was carefully placed to one side. A cut was made through the pelvic floor adjacent to the colorectum, and the entire length of the colon, from rectum to cecum, was excised and placed in ice cold PBS. Excised colon tissue was flushed with ice cold PBS to clear fecal contents, placed on bibulous paper, and cut longitudinally. Colons from non-tumor mice were cut in half longitudinally. One section was rolled, starting at the rectum, into a swiss roll and placed into saline-buffered formalin for 18-24 hours while rocking at room temperature, and then placed into 70% ethanol. The other half of the colon was scraped using a clean razor blade to harvest the epithelial contents and then snap frozen in an Eppendorf tube. For small intestine experiments, ~11cm of the ileum was harvested like the colon, swiss rolled, and fixed in saline-buffered formalin for 18-24 hours. The dissection of tumor mice after sacrifice was performed as above, but before fixation tumors >1mm were counted along the length of the colon (the rectum was not included). 1-2 tumors, greater than >2mm, were harvested from each mouse of available, combined in a single Eppendorf tube, and snap frozen. Statistical analysis for difference in gross tumor number was done using the Kruskal-Wallis test for multiple comparisons in PRISM.

Prior to preparation of colon and tumor tissue for western blot, RNA, and DNA analysis, tissue was pulverized under cryogenic conditions using a Bessman Tissue Pulverizer. For western blot analysis, a small amount of tissue was lysed in radioimmunoprecipitation (RIPA) buffer containing protease and phosphatase inhibitors, in combination cell disruption with 6-7 pumps of the cell solution through a 25-gauge needle. The lysis solution was then incubated at 4°C for 30 minutes with rotation and subsequently clarified 14,000 RPM in a chilled microcentrifuge. The supernatant protein concentration was then determined by the bicinchoninic acid (BCA) assay. Lysates were then equalized using RIPA buffer to 2-6 mg/m, diluted 5/6 using a 6x SDS-PAGE sample buffer and boiled for 10 minutes at 95°C. Protein lysates were stored at −80°C. 30-40µg of boiled protein solution was run a 12% polyacrylamide gel, and then transferred to nitrocellulose membrane using the semi-dry method (StandardSD program) provided by the Trans-Blot turbo transfer apparatus. Blots were blocked for 1 hour at room temperature using Intercept® tris buffered saline (TBS) blocking buffer, and subsequently transferred to Intercept® (TBS) antibody diluent solution containing primary antibodies for primary blotting overnight at 4°C. The following day, blots were washed three times using TBS containing 1% Tween-20 (TBST), blotted with secondary antibodies diluted in Intercept® (TBS) antibody diluent solution, and then washed four times with TBST. All secondary antibody steps were done at room temperature. Scanning of blots and quantification was done using a Li-Cor Odessey CLx. Primary antibodies used for western blotting of mouse tissues were from Cell Signaling Technologies and included pan-AKT (40D4) (Cat. #: sc-408; RRID: AB_630882), MEK1/2 (L38C12) (Cat. #: 4694; RRID: AB_10695868), p44/42 MAPK (ERK1/2) (L34F12) (Cat. #:4696; RRID: AB_390780), phospho-AKT (Ser473) XP® (Cat. #: 4060; RRID: AB_2315049), phospho-MEK1/2 (Ser217/221) (41G9) (Cat. #: 9154; RRID: AB_2138017), phospho-p44/42 MAPK (ERK1/2) (Thr202/Tyr204) (197G2) (Cat. #: 4377; RRID: AB_331775), and Vinculin (E1E9V) XP® (Cat. #: 13901; RRID: AB_2728768). Total RAS protein was detected in tissues via the RAS10 (Cat. #: 13901; RRID: AB_2728768) primary antibody from Millipore-Sigma. Secondary antibodies were obtained from Invitrogen and included Goat anti-mouse IgG (H+L) Highly Cross-adsorbed Secondary antibody, Alex Fluor 680 (Cat. #: A-21058; RRID: AB_2535724) and Goat anti-rabbit IgG (H+L) Highly Cross-adsorbed Secondary Antibody, Alexa Fluor Plus 800 (Cat. #: A-32735; RRID: AB_2633284).

Preparation of genomic DNA was performed using the DNeasy Blood & Tissue kit from QIAGEN per manufacturer’s instructions with an accompanying cell disruption of tissue in provided lysis buffer through a 25-gauge needle. Extracted gDNA was stored at −20°C. Genotyping experiments were done using PCR using the methods in the above section. Extraction of RNA was done using the RNeasy Plus kits from QIAGEN per manufacturer’s instruction. RNA was stored at −80°C.

### Detection of activated K-Ras protein in mouse tissue preparations

Detection of K-Ras protein activation was done using purified CRAF-RBD protein fragment fused to glutathione-S-transferase (GST-RAF-RBD) and expressed in the BL21 strain of *Escherichia coli* and purified as described previously (61). Pulldown of GTP bound Ras proteins from tissue lysates was done as described previously by our lab (13) and done the same day as lysis. Briefly, tissues were lysed in magnesium enriched lysis buffer (MLB), containing protease and phosphatase inhibitors, in combination cell disruption using a 25-gauge needle. The lysis solution was then incubated at 4°C for 30 minutes with rotation, and subsequently clarified 14,000 RPM in a chilled microcentrifuge. The supernatant protein concentration was then determined by BCA assay. The protein concentration was equalized using MLB and each pulldown consisted of 400µg of tissue lysate at a final MLB volume of 300µL containing 10µL of glutathione agarose loaded with GST-RAF-RBD. Binding between Ras and RBD was allowed to proceed for 2 hours at 4°C with rotation. The binding reaction was then stopped by washing agarose thrice with ice cold MLB. After the last wash, the 10 µL of 6x sample buffer was added to 10 µL of the washed agarose and boiled at 95°C for 10 minutes. K-Ras pulldown was assessed via western blot as described in the previous section using the RAS10 primary antibody (Cat. #: 13901; RRID: AB_2728768) and the secondary antibodies described above.

### Histology, immunohistochemistry, and immunofluorescence studies

Chilled paraffinized blocks were sectioned (5µm depth), and sections were affixed to glass slides by overnight incubation at 40°C. Slide-adhered tissue was deparaffinized by bathing slides in three rounds of HistoClear for 5 minutes per wash at room temperature, and then hydrated by two washes of 100% ethanol, one wash of 70% and 50% ethanol for 2 minutes each at room temperature. Antigen retrieval (e.g Ki67 at 1:200 dilution) for immunohistochemistry was performed using pre-boiled DAKO citrate buffer at pH 6.0 for 10 mins in the microwave (95-99C). The slides were cooled in running water, sections were encircled with PAP pen and washed twice with PBS to confirm integrity of the PAP pen. Next, slide-affixed tissues were carefully covered using DAKO EnVision+ peroxidase block for 15 minutes and rinsed twice using PBS. The sections were then blocked using DAKO serum-free protein block for 20 minutes. The serum-free block was removed, and the primary antibody diluted in DAKO antibody diluent were then carefully applied to the slide-affixed tissue using a micropipette, and incubation was allowed to proceed overnight at 4°C in a humid chamber. The following day, slides were rinsed three times with PBS, and then exposed to EnVision+ HRP labelled anti-rabbit polymer for 30 minutes at room temperature, and then rinsed three times with PBS. Slides were developed using 3’,3’-Diaminobenzidine (DAB) solution for 10 minutes and stopped by immersion in deionized water. A counterstain of hematoxylin was used for 1 minute, followed by immersion in deionized water and dehydrated through graded ethanol; once in 70% ethanol for 2 minutes, once in 95% ethanol for 2 minutes, twice in 100% ethanol for 2 minutes and three times in xylene for 5 minutes. Permount mounting medium was used to mount the slides and left to dry at room temperature for 24 hours before they were digitally scanned using Olympus BX-UCB.

For immunofluorescence studies, deparaffinization and antigen retrieval steps are performed as per IHC protocol. Sections were blocked using serum-free protein block and slides were washed with PBS. The two primary antibodies, Lysozyme (thermofisher, cat: RB372A, lot: 372A2207C) and E-cadherin (BD, cat: 610181, lot: 6315829), were diluted in DAKO antibody diluent at 1:200 and 1:750 respectively and allowed to incubate overnight at 4°C in the dark. The slides were washed three times with PBS and the secondary antibodies, Rabbit 488 (Alexaflour, ref: A11034) and MsIgG2a 594 (Alexafluor, ref: A21135), were applied at 1:500 dilution for one hour in a dark humid chamber. The slides were then carefully washed with PBS three times. To counterstain, Hochest 33342 (ref: C10337G, lot: 2730849) solution was diluted at 1:10,000 and applied to the slides. Following a 5 minute incubation period, the slides were washed with PBS, mounted with prolong diamond antifade mounting solution (Invitrogen, ref: P36970) and sealed with a coverslip (Avantor, ref: 28393-241). The slides were dried for an hour at room temperature and stored at 4°C until imaging (VS200 Olympus). Digitized versions of H&E, IHC and IF prepared slides were stored and analyzed using the OMERO (v5.22.1) Web portal.

Quantitation of crypt heights from colon swiss rolls was done essentially as describe by our lab previously (62), but with a few changes. In each colon swiss roll, every fifth crypt was analyzed starting at the rectum. If the crypt was whole (i.e. a visible inter-crypt space), the distance from the crypt base to the colon lumen was recorded, and if not, the location of the crypt was merely tagged. For statistical comparison of crypt lengths between mouse genotypes, data from each group were aggregated, and analyzed using the Kruskal-Wallis test for multiple comparisons in PRISM. To visualize crypt lengths along the colon axis, both measured and tagged crypts had to be located along the length of the colon, and this had to be done separately for the proximal and distal colons, as the proximal colon shows transverse folds which significantly alter the densities of crypts along the colon axis. Furthermore, we noted that the overall length of the colon could be affected by the presence of a K-Ras mutation. Thus, the distal and proximal were measured independently, as determined by the start of transverse folds in the proximal colon, and the distal colon was normalized 0-50% and the proximal colon 50-100%. Therefore, for the distal side of the colon, crypts were located along the colon axis with the following formula: **[1-((A-(A/B))/A)]*50**; where A=colon segment length and B=colon segment crypt #. For the proximal colon a similar formula was used: **[[1-((A-(A/B))/A)]*50]+50**. Each crypt measurement (or marker for crypt) was given a unique fractional distance from the rectum. Tabulated crypt measurements and distances from the rectum were combined for each experimental group, and then visualized in PRISM by fitting data to both a LOWESS (10-point smoothing window) spline and a smoothened (4 neighbor averaging) 2^nd^ order polynomial curve. ‘Marked crypts’ (i.e. any crypt measurements <25µm) were not included in graphic calculations. To calculate the proliferative zone in the colon swiss rolls, the Ki-67 positive zones, starting from the crypt base, were calculated for every fifth crypt. These data were not localized, rather aggregated per genotypic group and statistically analyzed using the Kruskal-Wallis test for multiple comparisons in PRISM. Counting of colon dysplasia frequency (i.e. pedunculated adenomas, sessile adenomas, and microadenomas) was determined on the basis of morphological changes in the H&E section and confirmed via Ki-67 staining.

### GTP hydrolysis and binding assays using NF1 protein

Human full-length proteins were prepared as in (13). The catalytic domain of human NF1 was prepared by the structural biology core of Dana Farber Cancer Institute. GTP hydrolysis by human KRAS, with and without the catalytic domain of NF1 was monitored using the GTPase-Glo assays from Promega. Reactions were done in a 384-well white walled plate. Conditions for a single well will be described. Intrinsic hydrolysis reactions consisted of 5µL of 2xGBN solution (20µM GTP and 0.5mM DTT in GTPase/GAP buffer supplied by kit) and 5µL of 2xRB solution (4µM KRAS protein and 0.5mM DTT in GTPase/GAP buffer supplied by kit). NF1 catalyzed hydrolysis reactions consisted of 5µL of 2xGBN solution (20µM GTP, 0.5mM DTT, and 2µM NF1 in GTPase/GAP buffer supplied by kit) and 5µL of 2xRB solution (4µM KRAS protein and 0.5mM DTT in GTPase/GAP buffer supplied by kit). GTP hydrolysis was allowed to proceed for 30 minutes at room temperature before addition of 10µL of reconstituted GTPase-Glo reagent. Conversion of remaining GTP to ATP was performed for 30 minutes at room temperature under constant agitation. After 30 minutes, 20µL of detection agent was supplied to the reaction well and after 5 minutes luminescence was measured using a Promega GloMax plate reader. For each condition, the experiment was repeated three times with five technical replicates each. The technical replicates were averaged, and data were analyzed statistically in the Prism (v10) using a paired t-test.

To test binding of KRAS protein to the catalytic domain of NF1, the 6xHis-tag on KRAS protein was cleaved using ProTEV Plus (Promega). At the same time, KRAS protein was pre-loaded GppNHp. The cleavage and nucleotide reactions were performed in 100µL at 30°C for 3 hours in the following formulations: 40µM of wildtype 6xHis-KRAS4B or its mutants G12D, A59T, or A59E, 2mM EDTA, 1mM DTT, 75U of ProTEV Plus, 5mg/mL of GppNHp, and 1X ProTEV buffer. The cleavage/exchange reactions were stopped by the addition of 4.8µM MgCl_2_, buffer exchange into stabilization buffer (20mM HEPES pH 7.5, 50mM NaCl, 5mM MgCl_2_, and 1mM DTT) using the 7MWCO Zeba Spin columns per manufacturers protocol. Next, remaining protease and un-cleaved KRAS was pulled out of solution using 10µL of HisPur^TM^ Cobalt resin (ThermoFisher) per protein solution under rotation for 1 hour at 4°C. These pullout reactions were spun down for 2 minutes at 4°C at 700xg and the top 85% of the supernatant was collected. Each solution was then diluted to bring the KRAS protein to 32µM, flash frozen, and stored at −80°C overnight. The following day, the cleaved KRAS4B proteins bound to GppNHp and at 32µM were evaluated for interaction with the 6xHis-tagged catalytic domain of NF1. Using stabilization buffer supplemented with 0.1mg/mL of GppNHp, a 110µL solution consisting of 10µL of HisPur^TM^ Cobalt resin, 2µM 6xHis-NF1333, 4µM KRAS4B protein, and 1mM DTT was created and allowed to gently mix via end-over-end rotation for 1-1.1 hours at room temperature. Each binding reaction was washed twice using 100µL of stabilization buffer supplemented with 0.1mg/mL of GppNHp and 2-minute centrifugation at 4°C and 700xg. Three samples were taken to examine the pulldown of KRAS-GppNHp by resin bound NF1, the original binding reaction, the second wash, and the final pulldown. Each sample was scaled to 50µL and boiled for 10 minutes at 95°C after the addition of 6x SDS-PAGE sample buffer. Two replicate pulldown experiments were performed, and proteins were analyzed by SDS-PAGE using either 26-well 12% polyacrylamide Tris-glycine gels from Bio-Rad, or 15-WedgeWell 4-20% Tris-glycine gradient gels from Novex.

### Bulk RNAseq of tumor tissues

Bulk RNAseq was done using RNA derived from tumor samples (above) and performed as previously described by our lab (63). Briefly, RNA concentration and quality was determined using an Agilent Bioanalyzer/TapeStation. 1 µg of RNA with a RIN score greater than 7 from each sample was used as input for RNA-Seq library preparation and poly-A enrichment was performed using the NEBNext Poly(A) mRNA Magnetic Isolation Module. The RNA-seq library was prepared using the NEBNext Ultra Directional RNA Library Prep kit from Illumina and quality control of the libraries was done using the TapeStation and single read sequencing of the libraries was done on a NextSeq 500 Illumina platform in a High Output mode for 75 cycles.

Quality control of RNA sequencing was using FastQC and Qualimap, and Salmon pseudo-aligner was used to map sequences and calculate transcript reads using the Ensembl mm10 mouse genome as reference. Differential gene expression was determined using DESeq2. Two separate datasets were collected and merged and underwent batch correction through DESeq2. Validation of batch correction was accomplished through principle component analysis (PCA) and UMAP. Gene Set Enrichment Analysis (GSEA) was performed on the merge and normalized dataset. The GSEA desktop application was used to run the analysis with the MsigDB Hallmark gene set on tumors of different genotypes. Default parameters were used and gene sets with nominal p-value < 0.05 and FDR q-value < 0.1 were considered significant. DE intersection analysis was used to identify differentially expressed genes that were found in the G12D and A59E and Nf1 knockout tumors.

### Generation of organoids from mouse colon epithelium and tumors

Derivation of mouse organoids from colon epithelium and tumors was performed in the following manner. Mice were sacrificed and dissected as above. For the colon epithelial organoids, the middle colon was excised and placed in 10 mL of ice-cold PBS with 1% PenStrep. The colon pieces were gently washed 3 times with PBS-PenStrep. After the third wash, 10mL of ice-cold solution PBS-PenStrep supplemented with 8mM EDTA was added to the excised colon strip and incubated at 4°C for 1.5-2 hours with gentle rotation. After incubation, the mixture was vigorously shaken for 30 seconds. Care was taken to resuspend the effervescent foam product of shaking, as the authors noted that dislodged crypts appeared to get trapped in this solution state. 10 µL of the colon solution was then examined under a microscope to determine the concentration of crypts/µL. A solution of 100-500 crypts was then spun down at 1000xg and resuspended in ice-cold Advanced DMEM/F12 culture medium, this solution was then spun down again, and crypts were suspended in Matrigel at a concentration of ~100crypts per 50 µL dome. After setting of the Matrigel, domes were bathed in 500 µL of colon organoid media.

Tumor organoids were cut from the mouse colon to avoid excessive epithelium and placed in 10 mL of ice-cold PBS-PenStrep. Tumors were washed with PBS-PenStrep and then placed in a PBS-PenStrep solution containing 2mM EDTA (Chelation buffer). Tumors were incubated for 1 hour on ice, without agitation, in the Chelation buffer. Tumors were then washed once with 5 mL of ice-cold chelation buffer, washed once with ice-cold PBS, and then suspended in 5 mL of ice-cold digestion buffer containing 84.4% advanced DMEM/F12, 2.5% FBS, 100 µg/mL of primocin, 400 U/mL of collagenase, and 12.5% Type II Dispase. Tumors were digested for 2 hours at 37°C with agitation at every 30 minutes. The tumor solution was then washed with 5 mL of PBS after centrifugation of cells at 200 xg for 3 minutes. Tumor cells were resuspended in 500 µL of PBS, counted, and then 10,000-15,000 cells were seeded into 50µL Matrigel domes. Once set, matrigel domes were bathed in 500 µL of colon tumor organoid media.

### Generation of Cre-inducible ERBB2 and EGFR Lentivirus

Flag-tagged ERBB2 (NM_001289938.2) gene fragment containing BamHI restriction sites were ordered from Azenta, PCR amplified using primer pairs Forward: 5’-CTCGGATCCATGGACTACAAGGACGACGATGACAAGGGTGGTTCTGGTAAGCTGCG GCTCCCTGCCAGT-3’, Reverse: 5’-AGTGGATCCTCACGAGTGGGTGCAGTTGA-3’). PCR products were cloned into the Zero Blunt TOPO (Life Technologies) vector, excised with BamHI restriction enzymes, and ligated into BamHI digested pTomo vector (Addgene catalogue number 26291) using T4 ligase overnight at 16°C. Insert orientation was validated by Sanger sequencing.

Efficiency of expression and Cre-specificity of cloned constructs was tested by transfection into HEK293T cells, in the presence or absence of CAG-Cre. HEK293T cells were grown to 50% confluence and transfected with ERBB2 pTomo constructs in the presence of CAG-Cre or empty vector control. 48 hours post-transfection, cells were lysed using 1 mM EDTA, 50 mM Tris-HCl, 420 mM NaCl, 0.5% NP40, pH 7.5 containing protease inhibitor cocktail (Sigma) (SBN buffer), incubated on ice for 30 minutes, centrifuged at 12,000xg for 15 min, and supernatants collected. Total protein quantification was performed using BCA protein assay. Equal amounts of protein (20 µg) were heated in NuPAGE LDS sample buffer and NuPAGE Reducing buffer at 80°C for 10 min, subjected to 10-15% SDS-PAGE, and transferred to nitrocellulose membrane using the Trans-Blot Turbo apparatus. Western blotting followed the protocol in the previous section. The primary antibodies Mouse anti-FLAG (Sigma #F1804) and anti-Cre (Novagen #69050) were used at 1:1000 dilution. The secondary antibodies Goat anti-Rabbit Alexa Fluor Plus 800 (Invitrogen #A32735) and Goat anti-Mouse Alexa Fluor 680 (Invitrogen #21058) were used in 1:5000 dilution

After validation, constructs were then packaged into Lentivirus (LV) using the following protocol. Constructs were transfected in HEK239T cells together with third generation LV vectors pMD2.G (Addgene #12259), pMDLg/pRRE (Addgene # 12251), pRSV-Rev (Addgene # 12253). Four days post-transfection, media was harvested, and viral particles were concentrated using Beckman ultracentrifuge, SW55 rotor 35,000 rpm, 1.5 hours at 4°C. Supernatant was discarded and pellets were stored overnight at 4°C and then frozen at −80°C for prolonged storage.

### Organoid growth experiments

Testing of colon epithelial organoid response to EGF and drugs was done using organoids derived from Fabp-Cre(+) mice that harbored either wildtype KRAS, KRAS G12D, or KRAS A59E. For both EGF and drug experiments, organoids were seeded at 800 cells/well in a 384-well plate. For the EGF response test, cells were mixed in a 20µL 10% slurry of GFR Matrigel and colon organoid media that either lacked EGF or contained 0.5µg/mL of EGF. At day 7, 2.2µL of PrestoBlue was carefully added to each well of the 384-well plate. After ~3 hours incubation at 37°C, cell viability was assessed by reduction of PrestoBlue and measured using a ClarioStar plate reader with excitation at 535-560 nm and detection of emission at 590-615 nm.

To test the colon organoid drug response, 364-well plates were setup in the same way as above, and Erlotinib, Trametinib and RMC4550 was dispensed using automated D300e Digital Dispenser (Tecan). Mouse tumor organoids isolated from Fabp-Cre(+); Apc(+/2lox14) mice harbored either wildtype KRAS, or KRAS G12D or KRAS A59E alleles for LV infection were recovered from Matrigel using Cell Recovery Solution (Sigma), trypsinized and centrifuged at 0.8g for 30 min with 30 mL LV (virus titer 1 x 10^8^ IFU/mL). Tumor organoids plated in 384 well plates (Corning) at a density of 1000 cells/well with 10% growth factor free (GFR) Matrigel (Corning #354230) and colon organoid media in a total volume of 20 mL per well. Erlotinib (or DMSO control) was dispensed using automated D300e Digital Dispenser. Murine cetuximab (mCetuximab) was created by the Wittrup lab at Massachusetts Institute of Technology (MIT) and detailed by Dr. Tisdale (39). For the experiments published here mCetuximab was generated in the HEK293T cells and purified via protein A resin and SEC by the Dana Farber Cancer Institute Structural Biology core (**Supplementary Fig. S5D**). For the drug experiments, the mCetuximab solution was too viscous for application to organoid Matrigel slurry to be applied robotically. Instead, mCetuximab was administered to 384-well plates manually.

For all drug experiments involving either colon or tumor organoids, drugs were administered the day after plating of cells (day 1). Furthermore, two 384-well plates were generated for each experimental group, to assess viability on day 1 and day 7 using CellTiter-Glo2.0 (Promega). The day 1 plate measurement was not treated with inhibitors. Normalization of drug response to cell growth rate was done using a published protocol (64). Data were plotted in PRISM.

## Acknowledgements

We would like to thank Dr. Lina Du and Dr. Michael Peluso from DF/HCC Mouse Engineering Core with their help in embryo injection of K-Ras(+/LSL-A59E) mES cells, and Sirano Dhe-Paganon of the Structural and Chemical Biology Center of DFCI. Furthermore, we would like to thank Dr. Debbie Burkhart for her careful reading of the manuscript during drafting.

## Table Legends

Supplementary table 1. Patient data for Ala59 mutant KRAS tumors

Supplementary table 2. Similarly dysregulated genes from KRAS G12D and KRAS A59E+NF1-null tumors.

## Supplementary Figure Legends

**Supplemental Figure 1.**
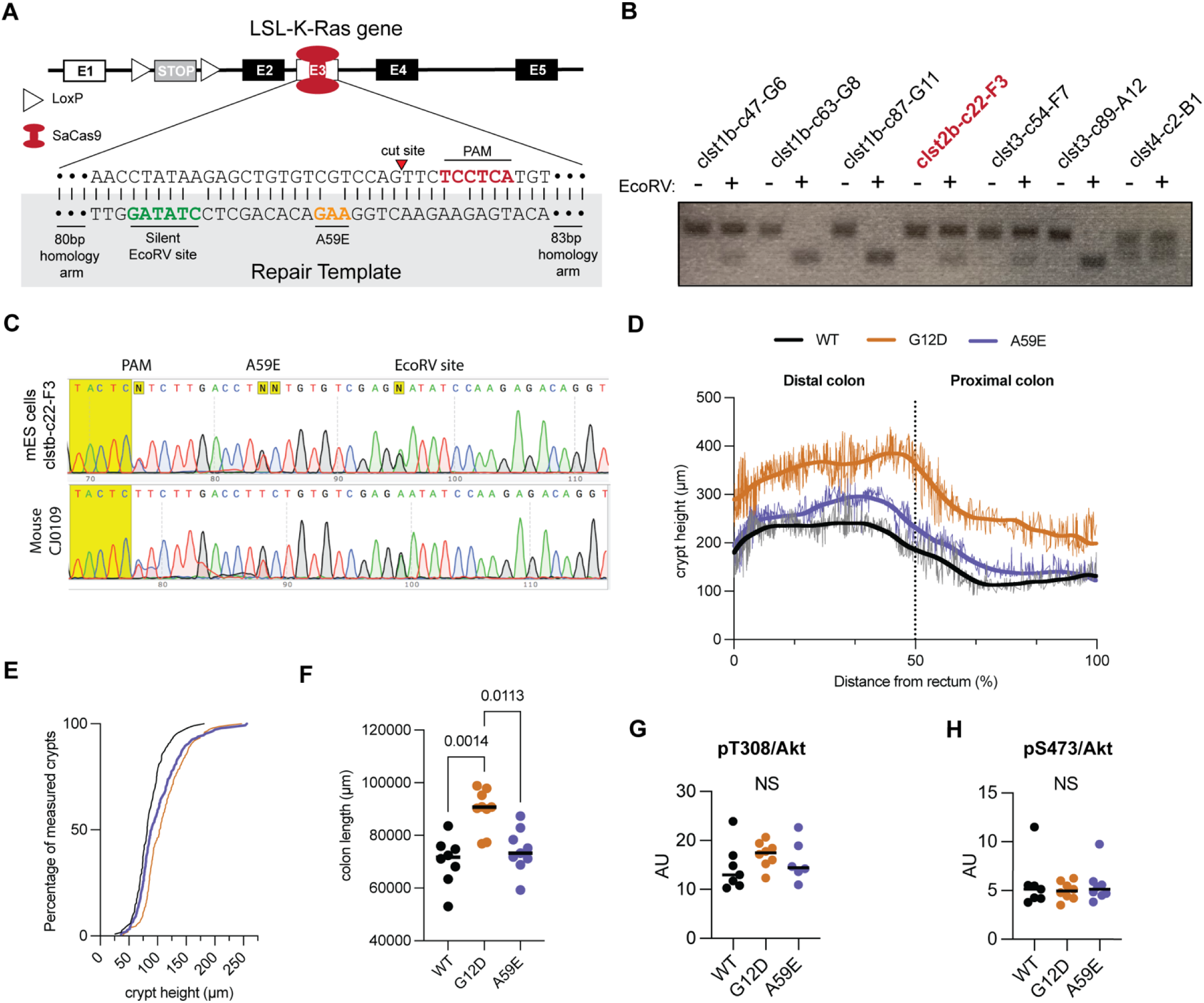
(A) Design of SaCas9 mutagenesis experiments to introduce the A59E mutation into LSL-K-Ras. Note that Exon 1 (E1) is non-coding. (B) 2% TAE agarose gel of selected mES cell clones harboring alterations at exon 3. EcoRV digest was used to detect the presence of silent mutation site (green base pairs in (A)). (C) Upper panel is exon 3 sequence of clone clst2b-c22-F3 and lower panel is representative F1 progeny showing germline transmission of *Kras^LSL-A59E/+^*. (D) Crypt heights by distance from the anus to the cecum. Thick line represents the calculated LOWESS line using a 10-point smoothing option. The thin jagged line represents a smoothened 2nd order polynomial calculated at every 10 points. Figure was generated in PRISM. (E) Frequency distribution of proliferating zone height measurements from Fig. 2D. (F) Length of colons from anus to cecum. (G,H) Phosphorylation status of Akt at Thr308 (G) and Ser473 (H) in total colon tissue. Statistical analysis of panels (F-H) was performed using Kruskal-Wallis tests with significance cutoff at <0.05.

**Supplemental Figure 2.**
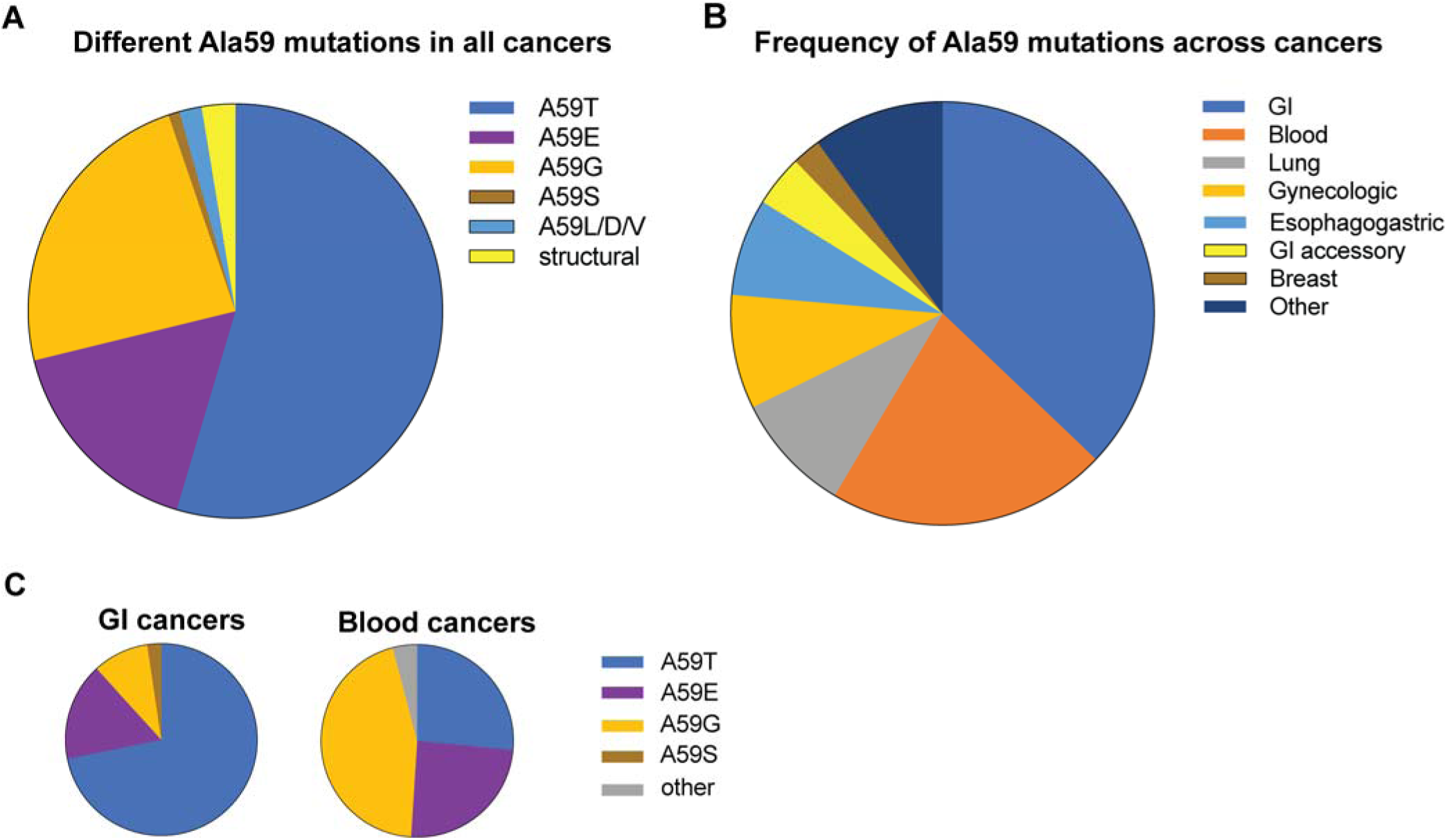
(A) Frequency of Ala59 mutants and variants across all cancers. ‘Structural’ variants refer to insertions, deletions, and frameshift alterations. (B) Compiled Ala59 mutant tumors (N=229). ‘Other’ designation includes Bladder, Brain, Kidney, Skin, and other cancers that occur at a frequency of <2.5%. See Supplementary Table 1. (C) Frequency of different Ala59 mutations for KRAS in GI cancers (N=84) and hematological cancers (N=45). ‘Other’ refers to single instance variants. See Supplementary Table 1.

**Supplemental Figure 3.**
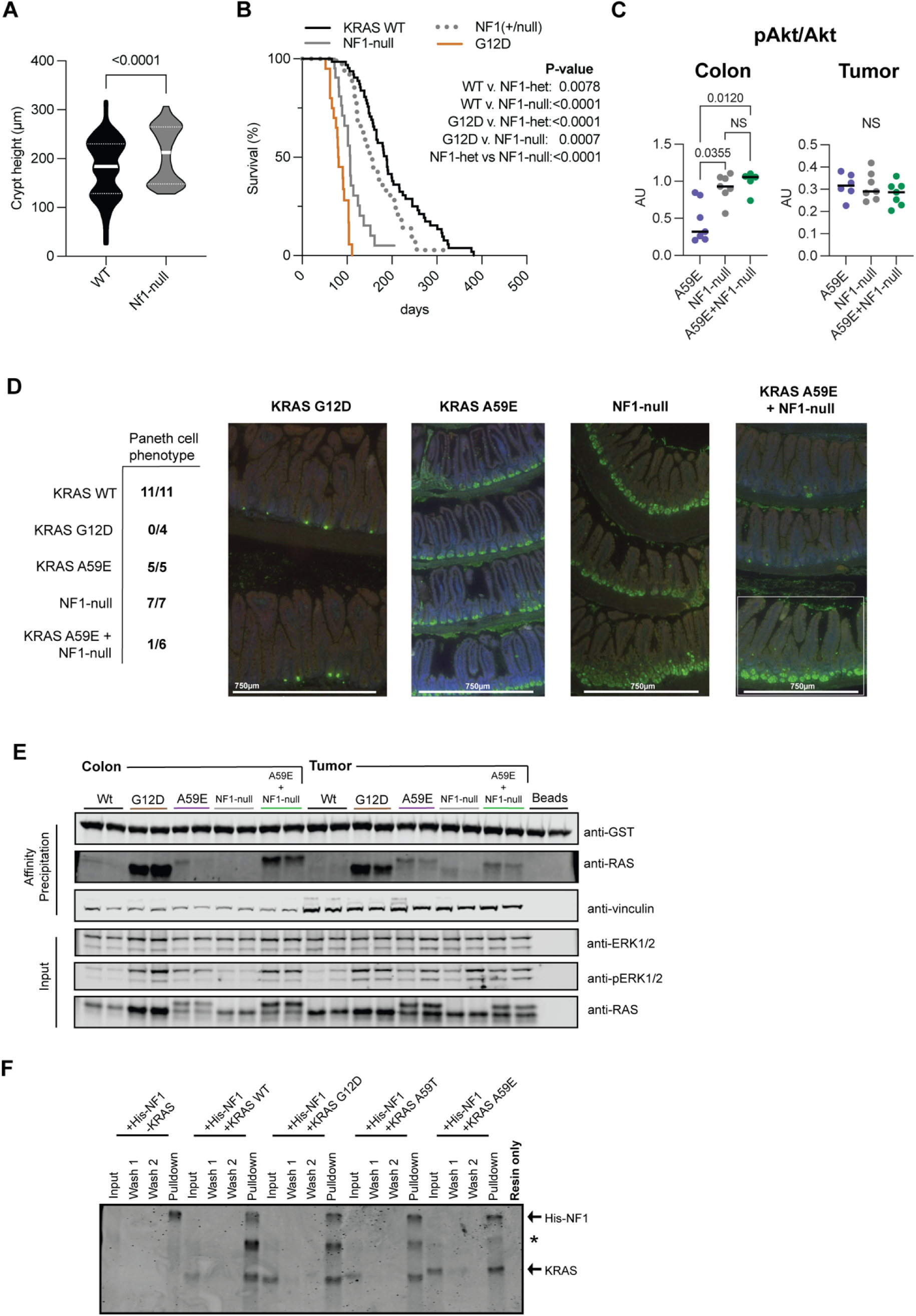
(A) Quantification of crypt lengths in Fabp-Cre(+) and Fabp-Cre(+), Nf1(fl/fl) mice. (B) Survival curve of Fabp-cre(+), Apc(+/2lox14) mice either wildtype for K-Ras, Nf1(+/fl), or were LSL-K-Ras G12D, Nf1-null, or harboring both KRAS mutation and NF1 loss. P-values are the product of Log-Rank Mantel Cox tests for survival. (C) Quantitative western blot analysis of AKT S473 phosphorylation from mouse tumors and colon scrapes. Statistical analysis of panels (C) was performed using Kruskal-Wallis tests with significance cutoff at <0.05. (D) Paneth cell staining of small intestines from Fabp-Cre(+) animals and the indicated genotypes. Paneth cells are indicated by lysozyme staining (green) at the bottom of the small intestinal crypts. Left, fraction of small intestines positive for Paneth cells. Right, representative images of small intestine examining immunofluorescence of lysozyme (green), E-cadherin (red), and DAPI (blue). Small intestine both positive and negative for lysozyme staining (right most panel) is from Fabp-Cre(+), K-Ras(+/LSL-A59E), Nf1(fl/fl) mice. Fabp-cre(+), K-Ras(+/+) mice images are not shown but are identical to Fabp-Cre(+), K-Ras(+/LSL-A59E) and Fabp-Cre(+), Nf1(fl/fl) mice. The white box shows the beginning of the jejunum which does not express Fabp-cre. (E) Left, western blot results from RAS activity assay using GST-CRAF-RBD from colon scrapes and tumors. Right, quantitation of KRAS A59E and G12D pulldown by GST-CRAF-RBD of Fabp-Cre(+), K-Ras(+/LSL-G12D) mice, Fabp-Cre(+), K-Ras(+/LSL-A59E) mice, and Fabp-Cre(+), K-Ras(+/LSL-A59E), Nf1(fl/fl) mice. Mice sacrificed for tumor tissues were all Apc(+/2lox14). (F) Repeat of experiment done in Fig. 3I. Pulldown of purified wildtype KRAS4B bound to GppNHp and its mutants G12D, A59T, and A59E by the His-tagged catalytic domain of NF1. * Indicates remaining un-cleaved 6xHis-KRAS4B.

**Supplemental Figure 4.**
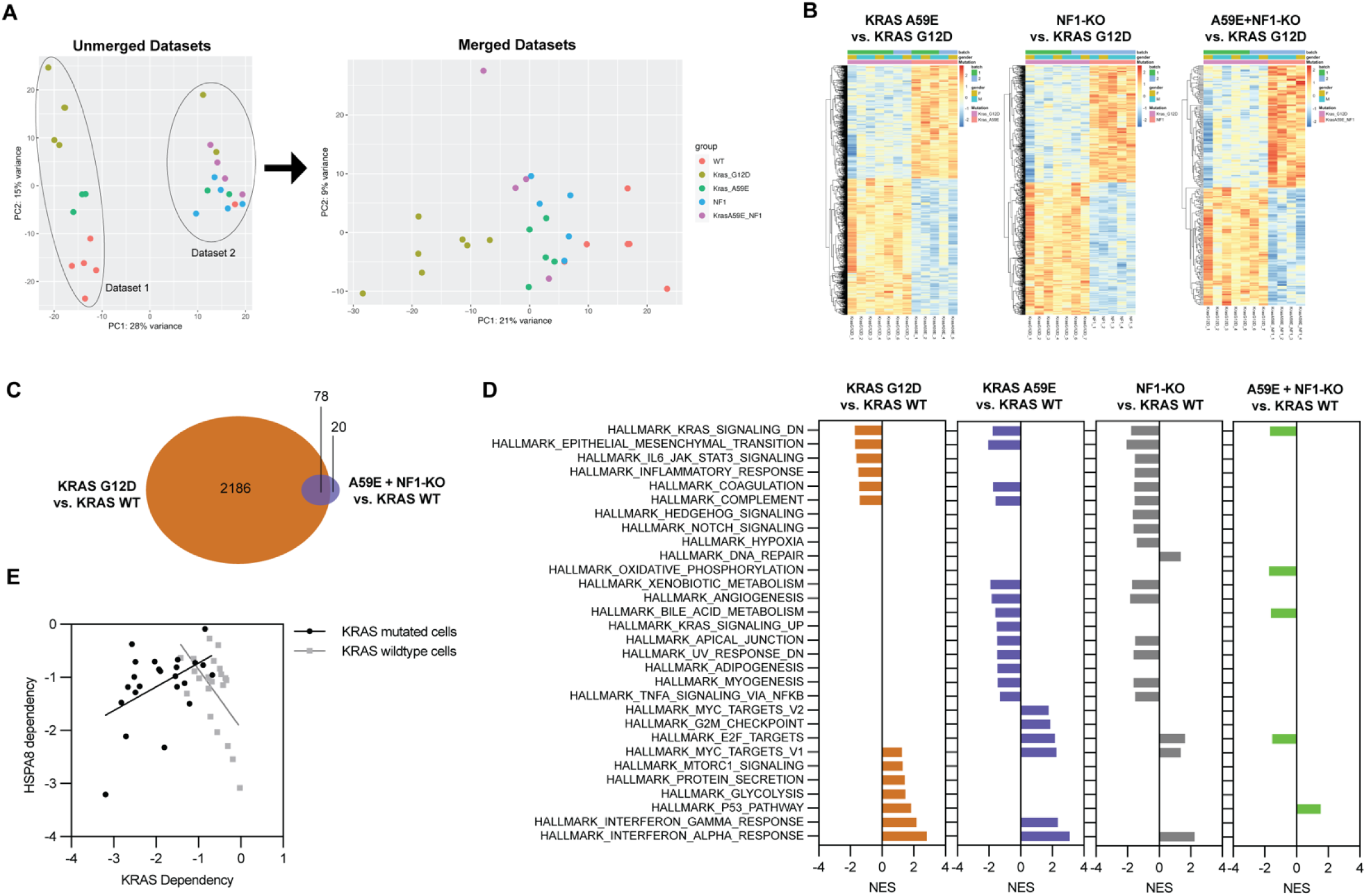
(A) PCA diagram showing data before and after merging and batch correlation. (B) Heatmap of differentially expressed genes (DEGs) in KRAS A59E, NF1-null, or doubly mutant tumors when compared to KRAS G12D. (C) Venn diagram comparing DEGs in KRAS G12D and KRAS A59E tumors in comparison to wildtype KRAS. (D) Gene set enrichment analyses of different mouse tumors harboring the indicated genotypes. Each dataset was compared to the same Fabp-Cre(+), Apc(-/-), K-Ras(+/+) dataset. (F) Correlation of HSPA8 and KRAS dependency scores of colon adenocarcinoma cell lines from DepMap. See Supplementary Table 2.

**Supplemental Figure 5.**
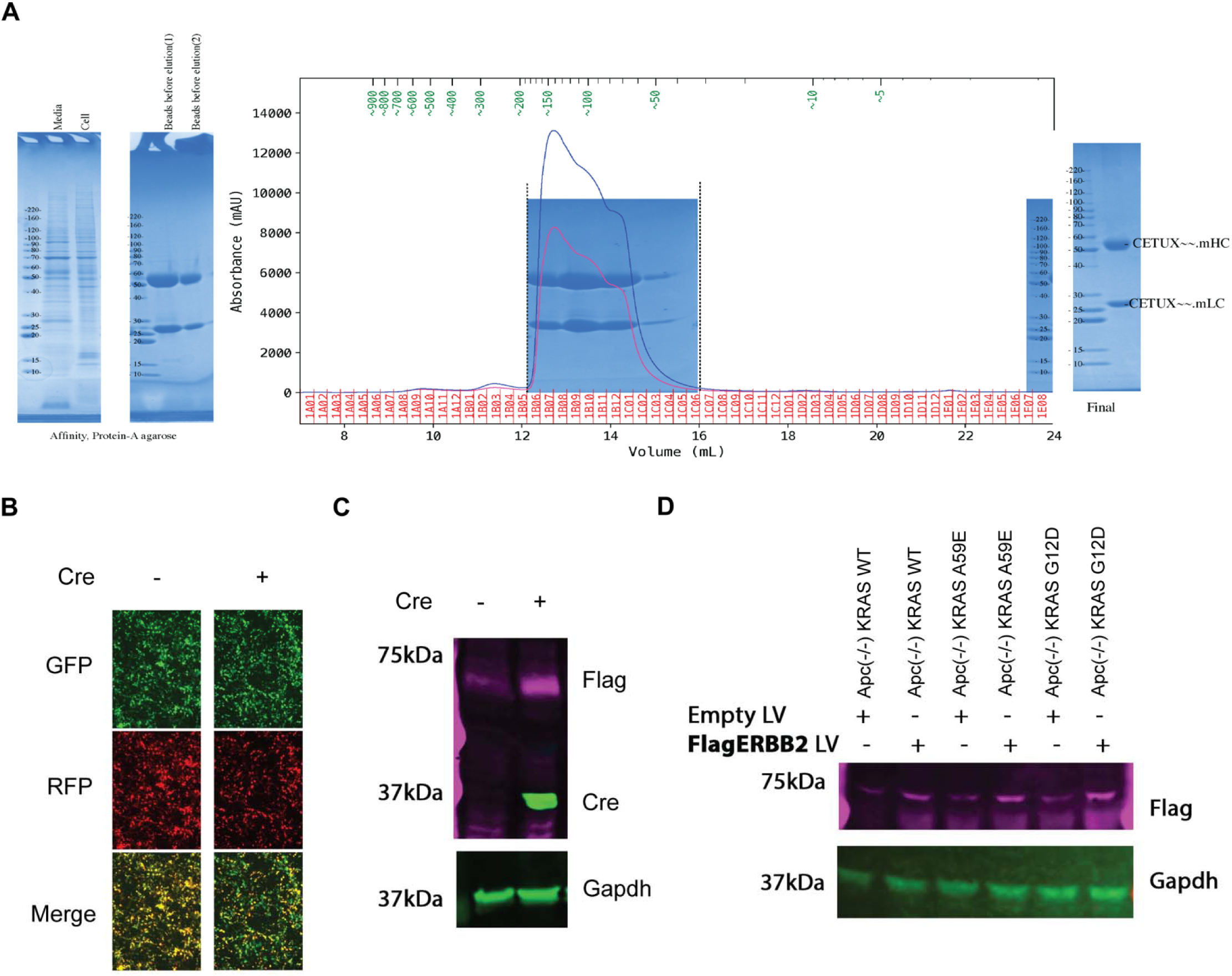
(A) Representative purification of mCetuximab generated from HEK293t cells. (B) and (C) Validation of pTOMO Flag-ERBB2 function in HEK293t cells after expression of Cre.

